# The mammalian plasma membrane is defined by transmembrane asymmetries in lipid unsaturation, leaflet packing, and protein shape

**DOI:** 10.1101/698837

**Authors:** JH Lorent, KR Levental, L Ganesan, G Rivera-Longsworth, E. Sezgin, MD Doktorova, E Lyman, I Levental

## Abstract

A fundamental feature of cellular plasma membranes (PM) is asymmetric lipid distribution between the bilayer leaflets. However, neither the detailed, comprehensive compositions of individual PM leaflets, nor how these contribute to structural membrane asymmetries have been defined. We report the distinct lipidomes and biophysical properties of both monolayers in living mammalian PMs. Phospholipid unsaturation is dramatically asymmetric, with the cytoplasmic leaflet being ∼2-fold more unsaturated than the exoplasmic. Atomistic simulations and spectroscopy of leaflet-selective fluorescent probes reveal that the outer PM leaflet is more packed and less diffusive than the inner leaflet, with this biophysical asymmetry maintained in the endocytic system. The structural asymmetry of the PM is reflected in asymmetric structures of protein transmembrane domains (TMD). These structural asymmetries are conserved throughout Eukaryota, suggesting fundamental cellular design principles.

## INTRODUCTION

The asymmetric distribution of lipids between the two leaflets of the plasma membrane (PM) bilayer is a prevalent and fundamental feature of cells across the tree of life ^1-3^. In mammals, compositional asymmetry is most widely studied for phosphatidylserine (PS), a negatively charged lipid found almost exclusively on the cytoplasmic (inner) leaflet of the PM. Other phospholipid headgroups are also asymmetrically distributed, with sphingolipids enriched in the exoplasmic leaflet, while charged and amino-phospholipids are more abundant on the cytoplasmic side ^2,4,5^. These distributions were originally established in red blood cells (RBCs) ^4,5^ and platelets ^6^ and later confirmed in PMs of various nucleated cell types ^1,7^. Asymmetric distributions were also proposed between the lumenal and cytoplasmic leaflets in organellar membranes ^8^.

Because of their reliance on chromatographic separate methods, previous measurements of lipid asymmetries in biomembranes have been limited to the few major lipid types defined by their polar headgroups. However, advances in lipidomics reveal a vast diversity of lipid species in mammalian membranes, comprising hundreds of lipid species with distinct headgroups, acyl chains, and backbone linkages ^9^. How these hundreds of individual lipid species are distributed across the PM, and there the detailed compositions of the individual leaflets, remains poorly understood.

Membrane functions are strictly dependent on biophysical properties – including membrane fluidity, permeability, lipid packing, intrinsic curvature, bending stiffness, surface charge, transbilayer stress profiles, and lateral domain formation ^10^ – which in turn are a consequence of membrane composition. Thus, one of the most important and challenging questions in membrane biology concerns the biophysical consequences of membrane asymmetry. While the connections between membrane composition and physical properties have been addressed extensively in studies of symmetric bilayers, little is known about how these insights translate to compositionally asymmetric membranes. Recently, robust protocols for producing asymmetric synthetic membranes ^11-14^ have been developed and used to characterize coupling (or lack thereof) across the bilayer ^15-17^, the effects of lipid asymmetry on protein conformation ^11,12^, and the influence of proteins on lipid distribution between leaflets ^18^. The extent to which membrane properties are coupled across the bilayer in living cells remains one of the foremost open questions in cell biology.

Disparities in biophysical properties between the two leaflets of the PM in living cells have long been proposed ^19^, but remain ambiguous due to experimental limitations and inconsistencies. Studies have often relied on RBCs, since they lack internal membranes, and exoplasmic quenchers or headgroup-specific chemistries to probe individual leaflets. However, even these seemingly simple methods yielded contradictory conclusions, with some studies inferring a more fluid inner PM leaflet ^20-23^ and others a more fluid outer leaflet ^24-29^. Ultimately, the presence and nature of biophysical asymmetry in mammalian membranes remains unresolved because of several limitations inherent to methodologies that rely on probes based on biological lipids and fluorescence quenchers (see Supplementary Text). Finally, previous studies have focused almost exclusively on the PM, which is amenable to analysis by virtue of being the most accessible membrane in the cell, and the only one in erythrocytes and platelets. Whether biophysical asymmetries are present in intracellular organelles has been rarely studied and remains unknown.

Here, we have investigated the lipidomes and biophysical properties of both leaflets in intact mammalian PMs. Using enzymatic digestion, the asymmetric distribution of ∼400 lipid species is defined for human RBC PMs and compiled into a detailed model for the compositions of PM leaflets. While the observed headgroup distributions are largely in line with previous reports, a striking asymmetry is observed for phospholipid acyl chains, with the cytoplasmic leaflet composed largely of highly unsaturated lipids. These observations suggested asymmetric properties in the two leaflets of the PM, an inference that was confirmed by atomistic simulations of complex lipid bilayers. These predictions were then directly examined using a novel approach that relies on selective staining of PM leaflets by a leaflet-selective, environment-sensitive fluorescent reporter. Clear differences in lipid packing are observed between the outer and inner leaflets of live cell PMs in two different nucleated cell lines and red blood cells, wherein a tightly packed exoplasmic leaflet apposes a more loosely packed cytoplasmic leaflet. These biophysical asymmetries persist in the membranes of intracellular endosomes. Finally, the asymmetric organization of mammalian plasma membranes is reflected in asymmetric structures of protein transmembrane domains, a property that appears to be conserved throughout Eukaryota, suggesting a conservation of lipid and protein organization in the membrane.

## RESULTS

### Detailed lipidomic asymmetry of a mammalian plasma membrane

The asymmetric distribution of phospholipid headgroups in eukaryote PMs was described by classical studies on mammalian red blood cells and platelets ^4-6^, and then confirmed in various mammalian cell types ^1,7^, as well as yeast ^30^ and nematodes ^31^. However, the differences in phospholipid backbones and acyl chains between exo- and cytoplasmic leaflets of the PM have not been investigated. Combining phospholipase digestion with mass spectrometric lipidomics, we report the detailed, comprehensive lipid asymmetry of the human red blood cell (RBC) plasma membrane. To this end, RBCs were treated with either phospholipases or sphingomyelinase to specifically digest only the lipid species present on the exoplasmic leaflet of the PM. These treatments were followed by quantitative mass spectrometry of ∼400 unique phospholipid species. The species remaining after enzymatic digestion are presumed to be on the inner leaflet (protected from the enzyme), whereas the enzymatically digested lipids are inferred to comprise the outer leaflet (Fig 1A and Supp Data). Combining several independent enzyme treatments allowed us to quantify the comprehensive lipidomes of both leaflets. The measured asymmetric lipid headgroup distributions (Fig 1A) were consistent with previous estimates ^4,5^. PLA2 treatment of intact cells minimally affected PE or PS, consistent with the near-absolute inner leaflet confinement of these aminophospholipids reported previously ^4-6^. PI, PA, and PE-O were similarly unaffected, suggesting their near-complete inner leaflet residence. SM was largely degraded (∼90%) by treatment of intact cells with SMase, confirming its concentration in the outer leaflet, while ∼60% of the PC was on the outer leaflet.

**Fig 1.**
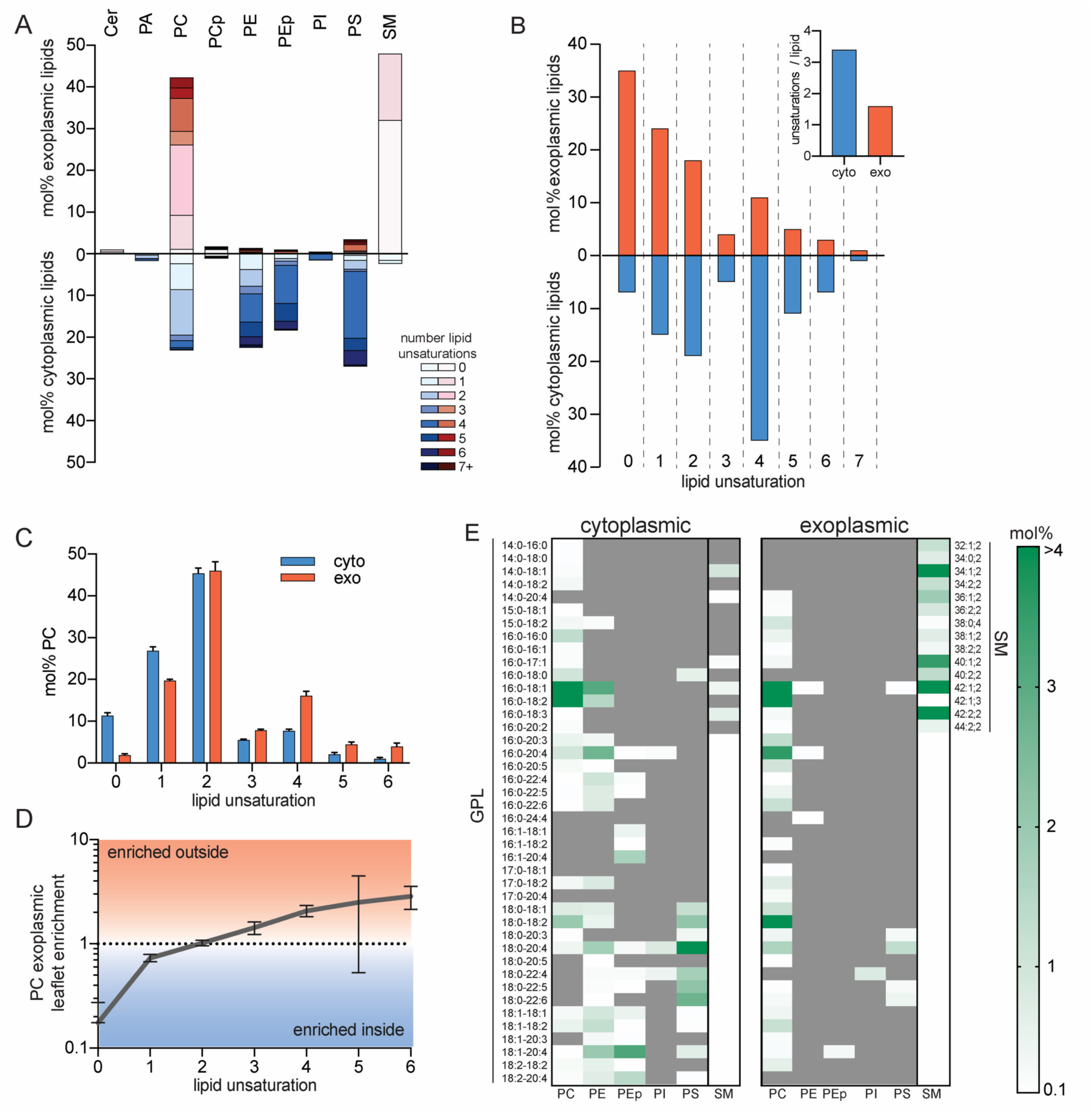
Lipidomic asymmetry of erythrocyte PMs. (A) Phospholipid compositions of exo- (red) and cytoplasmic (blue) PM leaflets as defined by enzymatic digestion and mass spectrometry. The exoplasmic leaflet is almost exclusively composed of PC and SM; the inner leaflet is approximately equimolar between PC, PE, PS, and PEp (plasmalogen). (B) Leaflet asymmetry of acyl chain unsaturation. The plurality of phospholipids in the outer leaflet are fully saturated, whereas the majority of the cytoplasmic leaflet is polyunsaturated. (inset) Abundance-weighted average unsaturation is ∼2-fold greater for inner leaflet phospholipids. (C) Asymmetry of acyl chain saturation for PC species. Fully saturated acyl chains are highly enriched in the inner leaflet PC and there is a (D) general correlation between outer leaflet enrichment of PC species and the extent of unsaturation. Data in C-D are mean ± SD of 7 independent samples. (E) Lipidomic bar codes of the inner and outer PM leaflets. Shown in white-green scale are all lipid species (GPL = glycerophospholipid, SM = sphingomyelin) comprising <0.1 mol% of lipids with mol% encoded in green intensity (darkest = most abundant). Gray are species below the 0.1 mol% threshold (including not detected).

The quantitative accuracy of these measurements is supported by several key controls: (1) complete digestion of target lipids in lysed cells (Fig S1); (2) lack of hemoglobin leakage in enzyme-treated cells (Fig S2); and (3) decrease in target lipids (e.g. SM) that quantitatively matches stoichiometric appearance of reaction products (e.g. Cer; Fig S1). The third point is crucial, as the target and product lipids use different standards and often different detection modes (e.g. positive versus negative ion) for quantification.

Analysis of lipid hydrophobic chains revealed a striking difference in acyl chain unsaturation between leaflets, with almost 40% of exoplasmic leaflet phospholipids being saturated (not including the sphingoid backbone double bond, which is a *trans* double bond near the headgroup and does not disrupt lipid packing) and the majority having less than 2 unsaturations per lipid (Fig 1B). In contrast, the cytoplasmic leaflet was highly enriched in lipids with polyunsaturated acyl chains, with the majority of lipids containing 4 or more unsaturations. Overall, the average outer leaflet lipid bears 1.6 unsaturations in comparison to 3.4 on the inner leaflet (Fig 1B, inset). Most of the lipid headgroup classes were confined to one of the PM leaflets, precluding any within-class asymmetry. The only exception present in both leaflets was PC, which showed a surprisingly asymmetric distribution: fully saturated PC species (e.g. dipalmitoyl PC) were almost exclusively present on the *cytosolic* leaflet, whereas exoplasmic PC showed a preference for polyunsaturated species (Fig 1C). More generally, there was a clear correlation between the unsaturation of PC species and their relative enrichment in the outer leaflet (Fig 1D). We speculate that this unexpected asymmetry of PC species may arise from fatty acid remodeling. In that scenario, the outer leaflet may better reflect *de novo* synthesized lipid species, whereas inner leaflets are remodeled by cytoplasmic machineries, consistent with observations of fatty acyl selectivity in these two pathways ^32^.

The complete phospholipid compositions (‘lipidomic barcode’) of the two leaflets are shown in Fig 1E (Supp Data) and summarized in Fig 2A&C. These data highlight several notable features: (1) the PM lipidome is dominated by ‘hybrid’ lipids (i.e. one saturated and one unsaturated acyl chain), with the major exception being highly saturated SM comprising ∼35% of outer leaflet PLs; (2) polyunsaturated lipids comprise two-thirds of inner leaflet PLs; (3) ∼33% of outer leaflet PLs are SMs bearing very-long chain (24:0 and 24:1) fatty acids. (4) lipids containing two unsaturated acyl chains are rare, comprising ∼10% of the inner leaflet and essentially not present on the outer leaflet; (5) ether linked lipids (i.e. plasmalogens) comprise a sizeable fraction (∼20%) of the inner leaflet; (6) there is clear headgroup selectivity for lipid acyl chains, with polyunsaturated fatty acids concentrated in PE and PS and wholly excluded from SM; (7) the major PC species on the outer leaflet bear di-unsaturated acyl chains.

**Fig 2.**
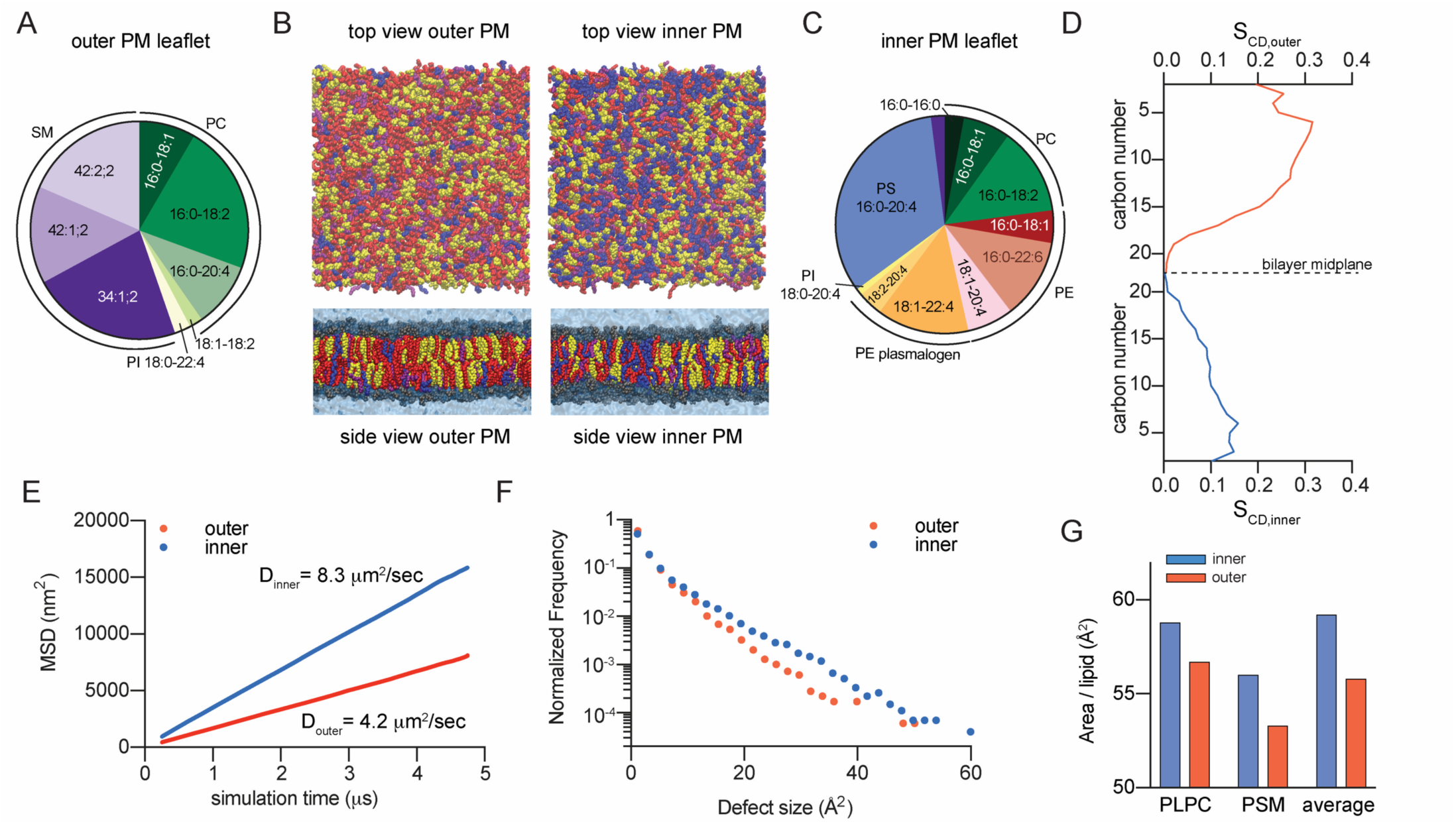
Atomistic simulation of biophysical asymmetry of erythrocytes PM. (A&C) Compiled compositions of inner (cytoplasmic) and outer (exoplasmic) PM leaflets from lipidomics. (B) Final snapshots of outer- and inner-PM leaflet mimetic simulations (yellow = cholesterol, red = saturated acyl chains, purple = mono- and di-unsaturated, blue = polyunsaturated; headgroups not shown in top views). (D) Concentration-weighted average order parameters for lipids in the outer versus inner leaflet simulation suggest a more ordered outer leaflet. (E) Slope of average MSD over time reveals ∼2-fold slower diffusivity of the simulated outer leaflet. (F) Histogram of hydrophobic defects reveals more abundant large defects in the simulated inner leaflet. (G) Area/phospholipid for PLPC and PSM (the two lipids shared between both simulations) and the abundance-weighted average phospholipid.

### Atomistic simulations demonstrate differences in leaflet physical properties

The striking differences in phospholipid composition between PM leaflets suggested the possibility of biophysical asymmetries across the PM. Differences in membrane properties associated with lipid headgroups (e.g. charge and size) have been extensively documented ^33-38^, but lipid acyl chains are also major determinants of membrane structure and organization ^10^. Namely, membranes rich in saturated lipids are more tightly packed, rigid, and ordered, whereas high unsaturation levels yield more fluid, loosely packed membranes, suggesting that the outer leaflet of mammalian PMs may be more tightly packed and ordered than the inner leaflet. This possibility was evaluated by atomistic molecular dynamics simulations ^39^. To specifically isolate the biophysical effects of distinct leaflet compositions from transbilayer asymmetry *per se*, we compared symmetric bilayers composed of inner- or outer-mimetic lipid complements (Table I). The full lipidomic complexity described in Fig 1 was distilled to a set of representative lipids for each leaflet that recapitulates the headgroup and acyl chain profiles of the compete leaflet lipidomes (Fig 2A&B; Table I; for details, see Supp Table II). Cholesterol was included in both systems at 40 mol%, since this was the abundance observed in RBCs overall, and cholesterol was found to distribute approximately equally between leaflets in previous PM-mimetic simulations ^40^ (cholesterol asymmetry is addressed in Discussion).

**Table I.** Distilled inner and outer leaflet lipidomes of the RBC PM.

| **(A) Outer (exoplasmic) leaflet** |  | **(B) Inner (cytoplasmic) leaflet** |  |
| --- | --- | --- | --- |
| **representative** | **mol% of** | **representative** | **mol% of** |
| PC 16:0-18:2 | 22.2 | PS 16:0-20:4 | 33.1 |
| SM 34:1;2 | 22.4 | PE O- 18:1-22:4 | 13.7 |
| SM 42:2;2 | 18.5 | PC 16:0-18:2 | 13.0 |
| SM 42:1;2 | 14.4 | PE 16:0-22:6 | 11.9 |
| PC 16:0-20:4 | 9.7 | PE 18:1-20:4 | 6.9 |
| PC 18:0-18:1 | 8.4 | PC 16:0-18:1 | 7.0 |
| PC 18:1-18:2 | 2.0 | PE 16:0-18:1 | 4.8 |
| PS 18:0-20:4 | 1.5 | PE O- 18:2-20:4 | 3.7 |
| PI 18:0-22:4 | 0.9 | PC 16:0-16:0 | 2.9 |
|  |  | SM 34:1;2 | 2.0 |
|  |  | PI 18:0-20:4 | 1.0 |

To quantitatively compare the two systems, we measured bulk parameters that reflect experimental measurements of membrane order, diffusivity, and headgroup packing. Overall lipid order was assessed by calculating the concentration-weighted average acyl chain order parameters (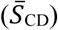 along both hydrophobic chains for all lipids in the systems (e.g. Fig S4). The outer leaflet-mimetic was roughly twice as ordered as the inner leaflet (Fig 2D), consistent with the inner leaflet being rich in relatively disordered unsaturated acyl chains. These differences were also reflected in the order parameters of identical lipids in the two mixtures, with 1-palmitoyl-2-linoleoyl PC (16:0/18:2 PC) and palmitoyl-SM being more ordered in the outer leaflet mixture (Fig S4). This order disparity corresponded to different molecular areas for various lipids comprising the inner versus outer leaflet simulations. The overall area per phospholipid (APL) was greater in the inner leaflet versus outer leaflet (59 versus 55 Å^2^), as also reflected in the APL for individual lipid species common to both simulations (Fig 2G). Finally, we observed striking differences in membrane fluidity, as the diffusion coefficient (D_leaflet_) (calculated from concentration-weighted mean square displacements (MSDs) of all lipids in the simulation) was ∼2-fold greater in the inner compared to the outer leaflet (Fig 2E). Finally, to explore differences in lipid packing at the headgroup level, hydrophobic packing defects were measured in the two systems. Such defects are regions of the membrane where hydrophobic acyl chains are transiently exposed to water^41^. The defect distribution is clearly shifted in the inner leaflet, indicating more abundant and larger packing defects, reflective of reduced headgroup packing (Fig 2F).

### PM leaflets have distinct biophysical properties

The atomistic simulations predict that membranes composed of inner leaflet lipids would have different physical properties from outer leaflet, as suggested by various experimental observations of broadly similar membranes. However, such insights have largely been based on investigations of symmetric membranes, as methodologies for robustly constructing asymmetric model membranes have only recently become widely accessible ^11-14,17,42,43^. Thus, the biophysical consequences of lipid asymmetry in either model or biological membranes remain poorly explored. The biophysical asymmetry of live cell PMs was probed using a fluorescent reporter of membrane packing (Di-4-ANEPPDHQ, Di4). The photophysical characteristics (e.g. fluorescence lifetime and emission spectra) of this dye are dependent on lipid packing ^44-49^, making it a robust reporter of membrane properties (Fig S5). The key feature making it suitable for leaflet-selective measurements is the presence of two charged moieties (Fig S6A) that prevent passive flip-flop across the bilayer ^45^. Thus, the exoplasmic leaflet of living cells can be selectively stained by adding the dye directly to the extracellular solution, whereas the inner leaflet is selectively stained by microinjecting the dye into the cytoplasm (schematized in Fig 3A). These features were confirmed in extensive control experiments described in the Supplement (Figs S6-S10). Specifically, we relied on back-extraction by BSA to assay the availability of Di4 to externally applied reagents. When Di4 was used to stain the outside of the cells (presumably resident on the outer leaflet), BSA extracted all PM fluorescence (Fig. S6B). On the other hand, BSA treatment did not affect PM fluorescence when Di4 was microinjected, confirming that it was inaccessible due to its residence on the inner PM leaflet. For completeness, we directly quantified the spontaneous Di4 flipping rate in synthetic membranes and observed minimal flip-flop on the experimentally relevant time scales (Fig. S6F), consistent with the structural features of this probe.

**Figure 3.**
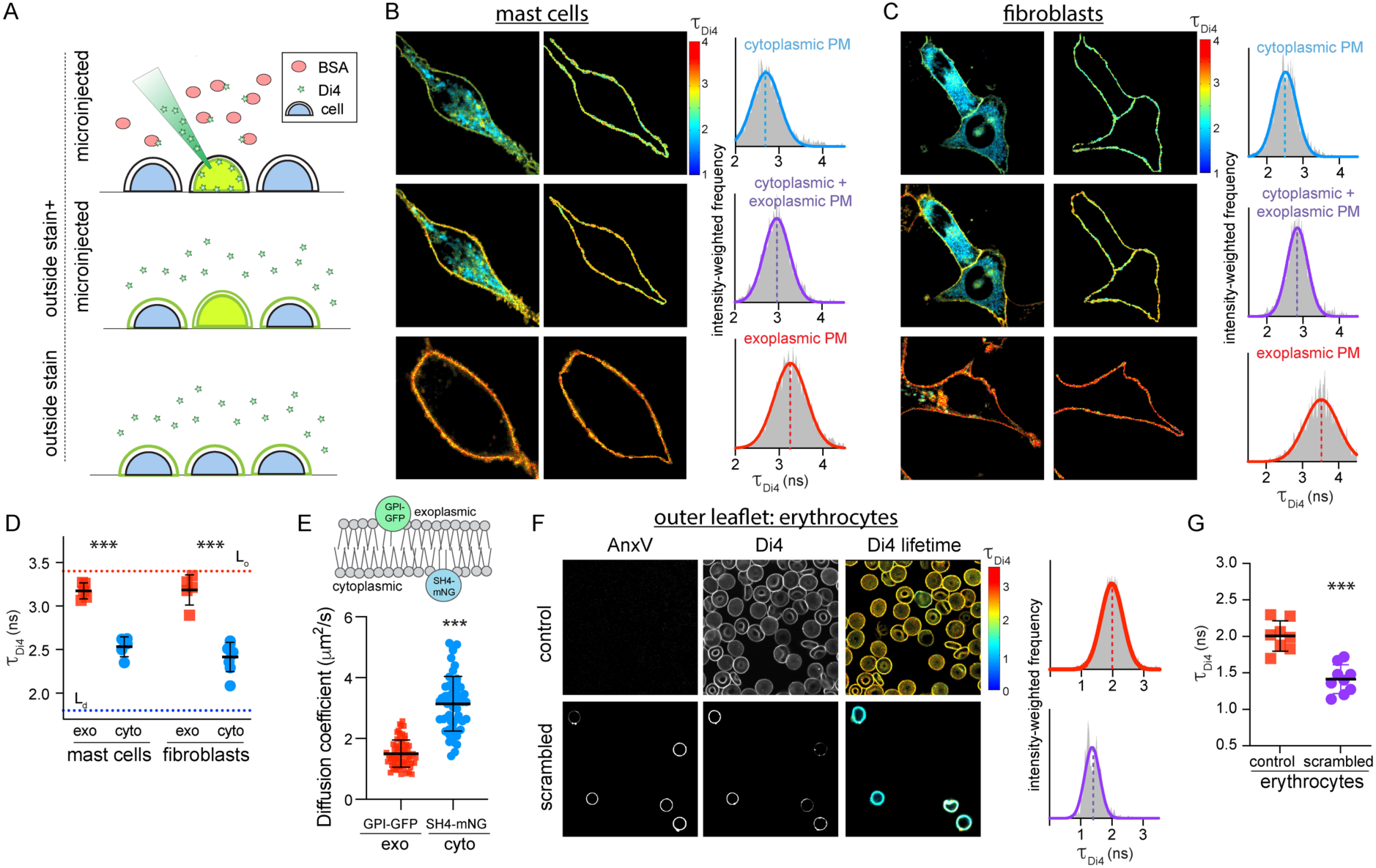
Biophysical asymmetry of the PM. (A) Microinjection of Di4 in presence of BSA stains only the cytoplasmic PM leaflet (microinjected). Subsequent addition of Di4 to the outside stains both membrane leaflets (outside stain + microinjected). Staining from the outside labels only the outer PM monolayer (outside stain). Exemplary FLIM images of (B) RBL mast cells and (C) 3T3 fibroblasts showing (left) whole cells, (middle) PM masks, and (right) intensity-weighted histograms of the PM mask. (D) Average Di4 lifetime in exoplasmic (red) versus cytoplasmic (blue) PM leaflets. Dotted lines represent Di4 lifetime in L_o_ and L_d_ phases in GUVs. Corroborating measurements of Di4 emission wavelength shifts are shown in Fig S9. (E) Diffusion coefficients measured by FCS of exoplasmically anchored GPI-GFP (red) versus cytoplasmically anchored SH4-mNG (blue) (F) AnxV staining for exoplasmic PS (left), Di4 intensity and lifetime (middle), and lifetime histogram (right), in control versus PMA-scrambled erythrocytes stained from the outside with Di4. PMA-induced scrambling induces PS exposure (AnxV binding) and reduces the packing of the outer leaflet. (G) Average Di4 lifetime in untreated (red) versus scrambled (purple) PM outer leaflets in erythrocytes. Data points in D and G represent averages of individual experiments, with 5-10 cells/experiment. Mean ± SD shown; ***p<0.001 for unpaired t-test. Points in E represent individual cells, with the mean ± SD shown; ***p<0.001 for unpaired t-test. All data are representative of >3 independent experiments.

Having confirmed Di4 as a leaflet-selective indicator of membrane physical properties, we probed their asymmetry in the PMs in two cultured mammalian cell types (RBLs and 3T3 fibroblasts). Note that while the lipidomic asymmetry was established in RBCs, we initially switched to nucleated cell types for the biophysical measurements because RBCs are not amenable to microinjection. In both cell types, microinjecting Di4 broadly stained intracellular membranes, with the PM easily identifiable both by the general morphology of the cell and the higher Di4 lifetime in the PM compared to internal membranes (left column of Fig 3B-C), revealing that the cytoplasmic leaflet of the PM is more tightly packed than those of intracellular membranes. An intensity-weighted histogram (right column) of a PM mask (middle column) revealed an average lifetime (τ_Di4_) of ∼2.5 ns in microinjected cells (Fig 3A-C, top row). When the same cells were then stained externally with Di4, the lifetime of the internal membranes was unaffected (Fig S7), whereas the Di4 lifetime in the PM increased significantly (Fig 3A-C, middle row), suggesting that the exoplasmic PM leaflet is more tightly packed than the cytoplasmic leaflet. This inference was confirmed by comparison to cells stained only from the outside, where Di4 signal was confined to the outer PM leaflet (absence of flipping prevents translocation to the cytosolic leaflet, while short incubations prevent significant endocytosis). In outside-only stained cells (Fig 3A-C, bottom row), Di4 lifetime in the PM was ∼3.2 ns, significantly greater than the PM in microinjected cells, revealing a striking asymmetry in membrane packing between the inner and outer leaflet. Similar values and trends were observed for both cell types (Fig 3D). To relate τ_Di4_ values to membrane properties, the cellular measurements were compared with those of L_o_ and L_d_ phases of synthetic phase-separated GUVs (Fig S5). The outer leaflets of cellular PMs were slightly less packed than the synthetic L_o_ phase, whereas the inner leaflets were approximately intermediate between the L_o_ and L_d_ phases. All FLIM results were corroborated by measuring Di4 emission wavelength shift, calculated as Generalized Polarization (GP, see Materials and Methods), in the same experimental setups (Fig. S9). To test the computational prediction that inner leaflet components diffuse faster than outer (Fig 2E), Fluorescence Correlation Spectroscopy (FCS) was used to evaluate the diffusion of fluorescent proteins anchored either to the outer leaflet (GPI-GFP) or inner leaflet (SH4 domain fused to mNeonGreen; SH4-mNG) by saturated fatty acids (Fig 3E). Outer leaflet-anchored GPI-GFP diffused ∼2-fold slower than inner leaflet SH4-mNG, in quantitative consistency with the computational result.

These measurements strongly suggest a biophysical asymmetry in mammalian PMs, with the outer leaflet being less fluid and more tightly packed. We confirmed that Di4 photophysical properties are not affected by charged lipids, pH, ionic composition of the media, or transmembrane potential (Fig S7), as partly previously described ^50^. Further, to rule out any artifacts associated with microinjection or the presence of internal membranes in nucleated cells, we investigated biophysical asymmetry in human erythrocyte PMs. These cells are too small for efficient microinjection, thus only the outer leaflet was probed by external staining. To assess biophysical asymmetry, asymmetric PMs in untreated RBCs were compared with those where asymmetry was pharmacologically abrogated. Specifically, treatment with phorbol myristate acetate (PMA) induces efficient PM scrambling, which is detected by the exposure of PS on the exoplasmic leaflet via the PS-binding probe Annexin V (AnxV) ^51^. This effect was confirmed, as untreated RBCs were almost exclusively AnxV-negative, whereas PMA-treated RBCs were uniformly AnxV-positive (Fig 3F, left column). Outer leaflet packing was significantly reduced by lipid scrambling, suggesting that the inner leaflet of the RBC PM is more loosely packed than the outer (Fig 3F-G). NR12S, an independent leaflet-selective order-sensitive probe, showed similar trends, ruling out probe-specific effects (Fig S10).

### Lipid packing asymmetry is preserved upon endocytosis

Having observed differential packing between the two PM leaflets, we next investigated whether biophysical asymmetries persist in endocytic membranes, which are composed in part of material arriving from the PM, but are also remodeled to comprise compartments for sorting, processing, and degradation ^52^. To probe the biophysical asymmetry in the endocytic system, these compartments were marked by pulse-chase with fluorescent dextran. Cells were incubated with fluorescent dextran for 2 h, during which the dextran is taken up by passive pinocytosis and accumulates in endocytic compartments, which were expanded by the increased load of non-degradable material (Fig 4A). The outer leaflets of cell PMs were then labelled by external Di4 (as in Fig 3), and the cells were incubated for up to 1 h (without Di4 or dextran) to allow dye endocytosis and staining of endolysosome membrane lumenal leaflets (Fig 4A). The contours of the dextran staining were then used as an endolysosomal membrane mask, which was overlaid onto the Di4 FLIM image (Fig 4A, mask). Strikingly, the Di4 lifetime in endocytic organelles was only slightly reduced compared to the outer PM leaflet (Fig 4). In contrast, the cytoplasmic leaflets of the dextran-positive endosomes probed by Di4 microinjection had a much lower membrane packing (i.e. τ_Di4_) than any of the lumenally stained endosomes. With longer incubation times, there was a slight reduction of τ_Di4_ in the lumenally labelled endosomes, likely due to membrane remodeling during endocytic maturation (Fig 4B) or to a slight degree of probe flipping between leaflets after long incubations (Fig S6F). These observations were completely consistent between cell types (RBLs in Fig 4; fibroblasts in Fig S11) and imply that biophysical membrane asymmetry is maintained in the endosomal system.

**Figure 4.**
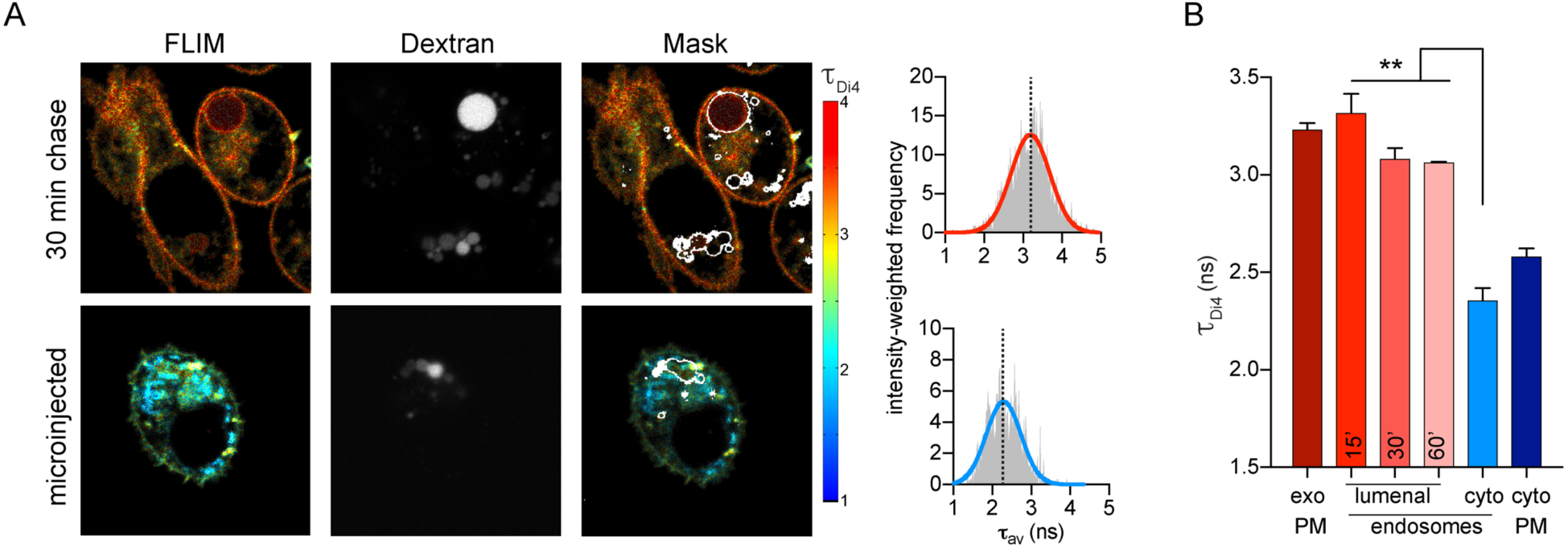
Asymmetry of membrane packing through the endocytic pathway. (A) Exemplary FLIM images of Di4 lifetime and dextran fluorescence in RBL cells following 30 min incubation to “chase” stains into endosomes. The accumulation of dextran (middle) is used to create an endosomal membrane mask (right images) to derive intensity-weighted histograms of τ_Di4_ (right). (B) The high lifetime (i.e. lipid packing) of the exoplasmic PM leaflet is maintained in the lumenal leaflets of endosomes, even up to 60 min after endocytosis. Images and quantifications of 3T3 cells are shown in Supp Fig S11. Mean ± SD of >10 individual cells; ***p<0.001 for unpaired t-test. Representative of at least 3 independent experiments.

### Structural asymmetry of transmembrane proteins conforms to lipid packing asymmetry and guides their trafficking

We recently reported that structural features of single-pass transmembrane domains (TMDs) determine protein partitioning between coexisting lipid phases in isolated plasma membrane vesicles ^53^. In particular, the lipid-accessible surface area of TMDs dictates partitioning to raft versus non-raft phases, with relatively thin TMDs (i.e. rich in Ala/Gly) more efficiently partitioning into tightly-packed raft-like domains, whereas larger TMDs (rich in Leu/Phe) are excluded from these regions. Our observations of packing asymmetries in live cell membranes prompt the hypothesis that proteins may have co-evolved to conform to these asymmetries. Namely, TMDs with relatively thin exoplasmic regions would minimally perturb the tight packing of the exoplasmic leaflet, whereas the TMD regions interfacing with the cytoplasmic leaflet could be larger due to their solvation by relatively loosely packed lipids. To evaluate this possibility, we calculated the ratio of exoplasmic to cytoplasmic TMD surface area for all annotated human single-pass PM proteins. This analysis revealed a clear asymmetry in TMD structures, with a distinct bias towards TMDs with relatively thin exoplasmic portions (Fig 5A), consistent with a previous report of an exoplasmic bias in the abundance of amino acids with smaller side chains ^19^.

**Fig 5.**
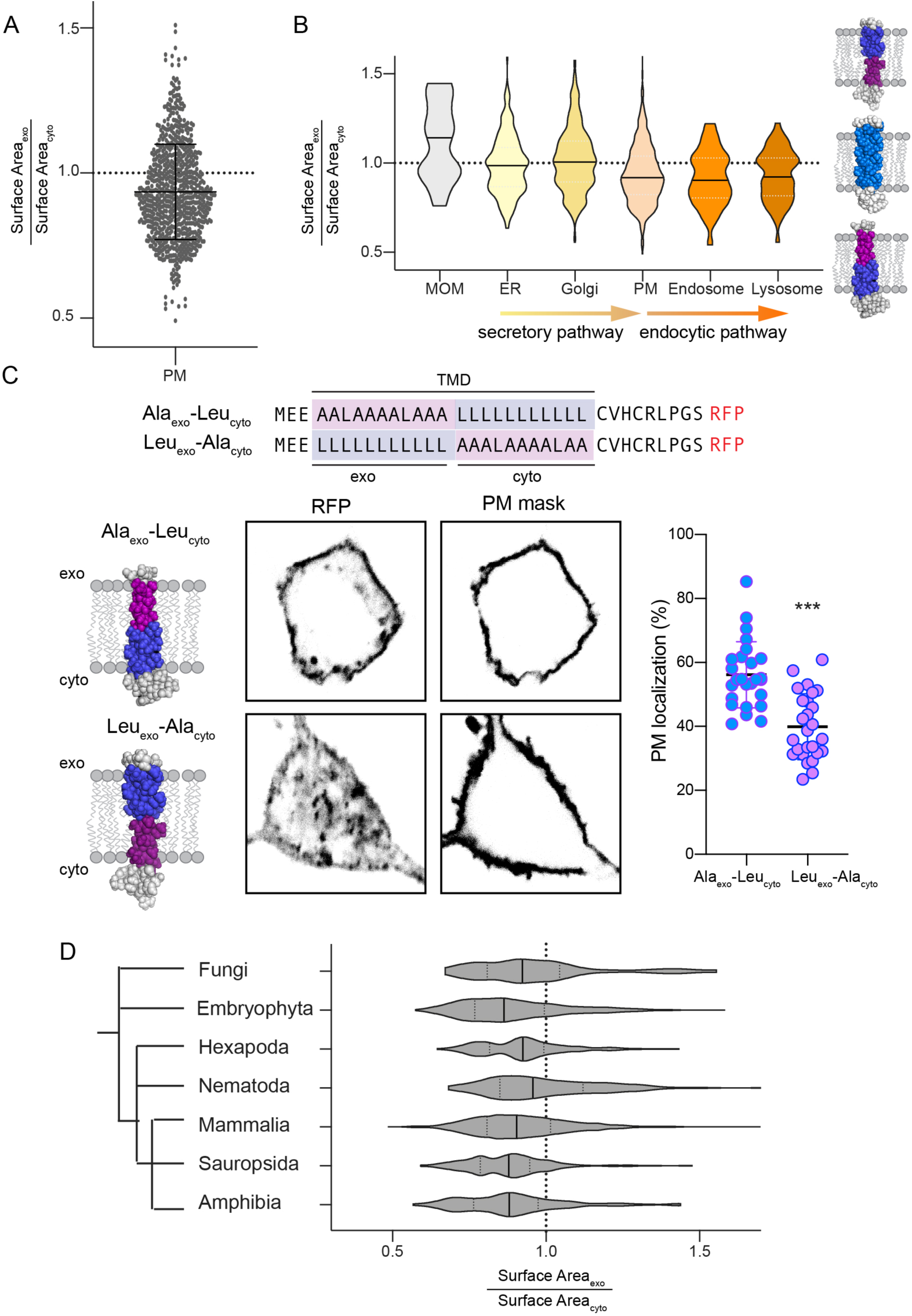
Structural asymmetry in PM protein TMDs is related to subcellular localization. (A) Asymmetry of single-pass transmembrane domain surface area between exoplasmic and cytoplasmic halves of human PM proteins. Shown are all annotated PM-resident, single-pass proteins in the human proteome (mean ± SD overlaid). (B) Violin plots demonstrating the distributions of relative exoplasmic / cytoplasmic surface areas for single-pass TMDs in various organelles. PM, endosomal, and lysosomal proteins are asymmetric with smaller exoplasmic halves, whereas ER and Golgi proteins are on average symmetric (MOM = mitochondrial outer membrane). (C) Subcellular localization of model TMDs that match (Ala_exo_-Leu_cyto_) or counter (Leu_exo_-Ala_cyto_) the biophysical asymmetry of the PM. The matching TMD localizes efficiently at the PM, whereas the countering TMD is largely intracellular. Mean ± SD of individual cells expressing the TMD construct; ***p<0.001 unpaired t-test. (D) Bias towards asymmetric TMDs is observed throughout eukaryote PMs. The phylogenetic dendrogram is intended to show only general evolutionary relationships.

The structural asymmetry of protein TMDs was not isolated to PMs but was also observed for proteins localized in endosomes and lysosomes (Fig 5B), consistent with the above observations of leaflet asymmetry in endosomes (Fig4A-B). In contrast, TMDs of the early secretory pathway, i.e. endoplasmic reticulum (ER) and Golgi apparatus, were on average symmetric (Fig 5B), suggesting that the biophysical asymmetry of the PM arises late in the secretory pathway. Interestingly, proteins of the outer mitochondrial membrane (OMM) have a clear bias towards the opposite asymmetry, with relatively small cytoplasmic regions. These analyses are fully consistent with our measurements of relatively tightly packed exoplasmic leaflets in the PM and endocytic organelles, suggesting that subcellular protein localization is determined to some extent by matching the physical properties of protein TMDs with their solvating lipids. To support this inference, we measured the subcellular distribution of model asymmetric TMDs designed to either conform to the measured biophysical asymmetry of the PM (i.e. thinner outer half; Ala_exo_-Leu_cyto_) or oppose it (thick outer half; Leu_exo_-Ala_cyto_) (Fig 5C). These TMDs are identical in all other “bulk” physical features (TMD length, surface area, hydrophobicity, etc). The majority (∼60%) of the asymmetric TMD that conforms to the packing asymmetry of the PM (thin outer half) localized to the PM, with a minor fraction in punctate endosome-like structures (Fig S13). In contrast, the TMD with the thicker outer half was largely confined to internal membranes (Fig 5C), identified as late endosomes via Rab7 staining (Fig S13).

Finally, the structural asymmetry observed in TMDs of mammalian PMs is also evident in non-mammalian eukaryotes. Across the eukaryotic domain of life, TMDs of single-pass PM proteins are highly asymmetric, with relatively thin exoplasmic regions and relatively thick cytoplasmic ones (Fig 5D). This finding suggests that the compositional and biophysical plasma membrane asymmetry we describe in cultured mammalian cell lines may be a fundamental and conserved design principle in Eukaryota.

## DISCUSSION

Combining enzymatic digestion with quantitative mass spectrometry revealed the detailed lipidomes of the inner and outer leaflets of RBC PMs. Consistent with expectations ^2,4,5^, SMs were highly enriched on the outer leaflet, whereas most glycerophospholipids were exclusive to the cytoplasmic leaflet, excepting PCs, which were distributed approximately equally between the two leaflets. Such significant lipid asymmetries are generally energetically unfavorable but can be maintained because of the intrinsically low rate of lipid flip-flop across the bilayer (typically estimated to be on the order of hours) ^54^. To establish and regulate asymmetry, cells rely on the coordinated action of numerous enzymes and channels. Enzymes called flippases and floppases move lipids between leaflets in an ATP-dependent fashion, while scramblases are channels that allow lipids to flow down concentration gradients ^54^.

It is important to point out that the reported compositions are not fully comprehensive. Complex glycolipids are not detected in the shotgun platform, nor are phosphorylated inositols. The most abundant glycolipid in human RBCs is globoside (Gb4), estimated to be <5% of all lipids ^55^; PIP_2_ is the most abundant phosphoinositide, estimated at <1% ^56^. Similarly, while the total concentration of cholesterol is quantified (∼40 mol%), its distribution between leaflets is not accessible by our methodology and remains controversial ^57^. Older measurements found conflicting results ^27,58^, and these contradictions are mirrored in more recent work, with some groups reporting inner leaflet enrichment of cholesterol ^59,60^ and others the opposite ^61^. Our measurements do not directly inform on this debate, though the higher packing of the outer leaflet may argue for higher cholesterol content therein. However, the packing of the inner leaflet is such that extremely low cholesterol concentrations are also unlikely. In the absence of consensus, and because cholesterol can rapidly flip between leaflets ^62-64^, the molecular dynamics simulations (Fig 2) assumed no *a priori* cholesterol preference for either leaflet.

The most notable disparity found between the acyl chains in the two RBC PM leaflets is ∼2-fold more double bonds per lipid in the inner versus outer leaflet (Fig 1B). This difference is likely responsible for the robust biophysical PM asymmetry we observe in RBCs and several nucleated mammalian cell types (Fig 3, Fig S9-10). These live cell results are consistent with model membrane observations suggesting that lipid order can be decoupled between the two leaflets of asymmetric membranes ^16,65^. The biophysical asymmetry appears to persist after internalization of the PM into the early endosomal pathway, suggesting that lipid asymmetry is also present in some intracellular compartments, consistent with other reports ^66^.

Comparing the lifetime of Di4 in the PM leaflets of live cells to synthetic model membranes suggests that the outer PM leaflet has biophysical properties similar to a L_o_ phase, whereas the inner PM leaflet is intermediate between the ordered and disordered phases (compare Fig 3 & S5). This comparison prompts examination of long-standing questions about the physical organization of the mammalian PM and its partitioning into ordered domains termed lipid rafts ^67^. Recent model membranes experiments revealed that ordered domains in phase-separating leaflets can induce domains in apposed leaflets which otherwise would not form L_o_ phases ^42,68,69^. Long-chain SM species are implicated as central mediators of such transbilayer domain coupling ^42^. The asymmetric RBC lipidome reveals an outer leaflet rich in high-melting lipids (i.e. saturated SM) and cholesterol, with a non-trivial abundance of low melting lipids (i.e. unsaturated glycerophospholipids). Such a membrane monolayer is poised to form coexisting ordered and disordered domains ^70-72^, with an area fraction dominated by the ordered phase. The inner leaflet contains largely low-melting unsaturated lipids that are not amenable to ordered domain formation, though the high abundance of long-chain SM species in the outer leaflet may promote domain coupling between leaflets. The ultimate organization of the membrane is then a combination of these membrane-intrinsic effects and extrinsic inputs like protein scaffolds ^73^ and cytoskeletal dynamics ^74,75^.

The lipid packing asymmetry of the mammalian PM is reflected in the structure of its resident transmembrane protein domains. We have recently shown that tightly packed membrane phases tend to select TMDs based on their lipid-accessible surface area, with larger, bulky TMDs being excluded in favor of thin TMDs ^53^. Consistent with these general principles, mammalian PM proteins have asymmetric surface areas, with the TMD region in the exoplasmic leaflet of the bilayer being relatively thin. Similar tendencies have been previously reported in bioinformatic studies of both mammalian and yeast PM proteins ^19^, with the authors anticipating their mechanistic origins as being the biophysical asymmetry reported here.

The TMD asymmetry of PM proteins is also evident in resident proteins of the late secretory and endocytic systems, consistent with our findings of asymmetric biophysical properties in endocytic organelles (Fig 4). In contrast, transmembrane proteins of other organelles, including the ER and mitochondria, have relatively symmetric TMDs, serving as an important control and also suggesting that these membranes may not be asymmetric. The physical characteristics of protein transmembrane domains have been previously implicated in subcellular protein sorting ^19,52,53^, with longer and thinner TMDs preferentially trafficking to the thicker, more tightly packed PM. A similar ‘matching’ between transbilayer membrane packing profiles and TMD structural appears to also affect PM localization (Fig 5C), as model TMDs conforming to the structural asymmetry of the PM are trafficked efficiently to the PM, whereas those with the opposite asymmetry are retained in intracellular membranes. Mechanistically, these observations suggest that sub-domains of intracellular sorting organelles (e.g. trans-Golgi and endosomes) select proteins with specific properties for transport to the PM. Alternatively, PM proteins that do not conform are preferentially endocytosed.

The inference that the structural asymmetry of PM TMDs is ubiquitous in eukaryotic organisms prompts the conclusion that biophysical PM asymmetry is a conserved and fundamental architectural principle of eukaryotic membranes. This possibility raises a key question: what adaptive advantage is gained from PM asymmetry? The generation and maintenance of lipid disparities in the two leaflets is likely energetically costly, suggesting a significant benefit to maintaining their highly asymmetric distribution. There are several hypothetical explanations for the benefits of PM asymmetry: from the standpoint of material properties, coupling distinct leaflets may combine desirable features into a single bilayer. For example, a tightly packed outer leaflet may serve as an effective permeability barrier, while the more fluid inner leaflet allows for rapid signal transmission. It may be interesting to develop methods to individually tune leaflet compositions and physical properties to determine whether certain cellular functions (e.g. signal transduction) are more dependent on one leaflet versus the other. A non-exclusive alternative is that PM asymmetry is used for energy storage, in analogy with energy storage by pumped-storage hydroelectricity. Cells may store the potential energy generated by pumping lipids against their concentration gradients, to be released later upon regulated scrambling of the bilayer. Finally, membrane asymmetries could be used as organellar identifiers for selective sorting of lipids and proteins, both between organelles and within the PM itself. In that context, the magnitude and mechanisms of transbilayer coupling between disparate leaflets in asymmetric membranes is an essential next step. Finally, it has been shown that membrane asymmetry can affect the properties of membrane domains ^14,17,42^ and possibly the partitioning of proteins between ordered and disordered lipid environments ^76,77^. Ultimately, deciphering the purpose of membrane asymmetry is central to understanding the functions of cell membranes.

## List of Supplementary Materials

Materials and Methods

Figure S1-S13

Supplementary Discussion

Tables S1 and S2

Supplementary References

Supplementary Data – Asymmetric lipidomes

## Acknowledgements

All fluorescence microscopy was performed at the Center for Advanced Microscopy, Department of Integrative Biology & Pharmacology at McGovern Medical School, UTHealth. We gratefully acknowledge Kai Simons, Theodore Steck, Yvonne Lange, and Gerald Feigenson for their critical feedback on this manuscript. Funding for this work was provided by the NIH/National Institute of General Medical Sciences (GM114282, GM124072, GM120351), the Volkswagen Foundation (grant 93091), and the Human Frontiers Science Program (RGP0059/2019). ES is funded by Newton-Katip Çelebi Institutional Links grant (352333122). Anton2 computer time was provided by the National Resource for Biomedical Supercomputing (NRBSC), the Pittsburgh Supercomputing Center (PSC), and the Biomedical Technology Research Center for Multiscale Modeling of Biological Systems through grant P41GM103712-S1 from the National Institutes of Health. All authors have no competing interests.

## Author contributions

JHL, IL, EL, and KRL designed the study. JHL, KRL, LG, GR, MD, and ES performed experiments. EL performed and analyzed the molecular dynamics simulations. JHL did the bioinformatics analysis. JHL, KRL, and IL analyzed the experimental results and wrote the paper. None of the authors have competing interests.

