## Supplementary Methods and Figs for "The mammalian plasma membrane is defined by transmembrane asymmetries in lipid unsaturation, leaflet packing, and protein shape"

### CONTENTS:

Materials and Methods
13 supplementary figures
2 tables
Supplementary Discussion

### MATERIALS AND METHODS

#### *General experimental design*

Lipidomics: Freshly isolated, intact human erythrocytes were treated phospholipase A2 or sphingomyelinase to specifically digest only the lipid species present on the exoplasmic leaflet of the PM. Comparison of enzyme-treated cells to untreated controls revealed the extent of digestion (i.e. abundance on the exoplasmic leaflet) of each of the ~400 unique phospholipid species detected in our measurements. Detailed asymmetry was measured for nearly all major phospholipid species in RBCs, including all PC, PE, PI, PS, SM, and the ether form of PE (plasmalogen; PE-O). The only exception was the PC plasmalogen (PC-O), which were not susceptible to any of our enzyme treatments. These lipids comprise less >1% of the total lipidomes and were assumed to distribute like other PC lipids. For all enzyme treatments, hemolysis was monitored to ensure that RBCs remained intact (Fig S2). Treatment of sonically disrupted cells ensured that the enzyme conditions were sufficient to completely degrade all available target lipids, with the increase in degradation products quantitatively validating the enzymatic digestions (Fig S1).

Spontaneous phospholipid flipping is negligible on our experimental time scales [1]. PLA2 treatment of intact cells did not degrade PE or PS to any appreciable extent, consistent with the near-absolute inner leaflet confinement of these aminophospholipids noted previously. This result further confirmed that minimal flipping of inner leaflet lipids occurred during our procedure. The observed headgroup distributions are in excellent quantitative correspondence with previous reports[2-4].

The lipidome of the inner leaflet was inferred from the lipid species remaining after digestion of intact cells. Conversely, the outer leaflet was calculated by subtracting the abundance of each lipid species remaining after digestion from the untreated controls. These data were highly consistent between multiple healthy adult donors (N=3) and repeated samples from the same donor (Fig S1). These measurements produced the raw lipidomes of the inner and outer leaflets given in Supplementary Data. These were then collated by sorting all species above a threshold of 0.5 mol% into self-similar groups defined by headgroup, acyl chains, and compositions (Table I and Supp Table II). The details for these groupings are in the legend to Supp Table II. These collated lipidomes largely recapitulate the headgroup and acyl chain profiles of the complete leaflet lipidomes.

Leaflet selective biophysical properties: The biophysical asymmetry of live cell PMs was probed using a fluorescent reporter of membrane packing (Di-4-ANEPPDHQ, Di4) (Fig S5A). To calibrate the sensitivity of Di4 to membrane packing, we constructed synthetic Giant Unilamellar Vesicles (GUVs) composed of DOPC, DPPC, and cholesterol. Such vesicles separate into microscopic liquid-ordered ($L_o$) and liquid-disordered ($L_d$) domains that have been widely employed as synthetic analogues of membrane structures in mammalian cells [5-7]. Di4 can be used to probe physical differences between phases because its emission spectrum is dependent on membrane packing, red-shifting in a more disordered environment, as typically described by the Generalized Polarization (GP) [8-10]. We observed a clear difference in GP between coexisting phases in our GUVs (Fig S5). The difference between $L_o$ and $L_d$ phases was much better resolved by measuring Di4 emission lifetime using Fluorescence Lifetime Imaging Microscopy (FLIM), as previously noted [11]. Fig S4 shows that average lifetime histograms in the two phases are distinct and non-overlapping, as opposed to those of GP in the same vesicle.

Having confirmed that Di4 sensitively reports membrane physical properties, selectivity for staining individual PM leaflets in live cells was verified. Cells were first stained externally by adding dye directly into the medium. The charged groups on Di4 should prevent flipping across the bilayer midplane, thus only staining the outer leaflets of the PM. This supposition was supported by BSA back-extraction experiments, wherein BSA complexes and removes any accessible dye from the exoplasmic leaflet of cellular membranes [12]. Application of BSA to externally stained cells completely eliminated Di4 fluorescence specifically at the PM, leaving only minimal residual signal from internal membranes (Fig S6B). This effect was observable even after relatively long incubation times: after 30 min of incubation, some dye was efficiently internalized into punctate endocytic structures (Fig S6B). Treatment with BSA did not noticeably affect the intensity of these structures, but did eliminate all observable signal from the PM, suggesting that Di4 does not flip across the bilayer in live cells on our experimental time-scales. These effects were completely independent of cell type.

To selectively stain the inner leaflet of the PM, cells were microinjected with Di4, together with a fluorescent dextran to ensure membrane integrity of the injected cells (Fig S6C). Due to leakage of dye from the micropipette, cells in the vicinity of the injected cell were also stained, though presumably only on the outer leaflet of the PM, as no internal membrane staining was observed (Fig S6C), suggesting the dye was unable to passively diffuse through the PM. Inclusion of BSA into the extracellular medium captures the leaked dye from the micropipette, preventing staining of nearby cells (Fig S6D and S7); however, the microinjected cells were still efficiently labeled on cytoplasmic membranes, including clear staining of their PMs (see Fig 3B-C, Fig S6D, and Fig S7A). This PM staining is exclusive to the cytosolic leaflet, as extracellular BSA did not eliminate this PM signal, in contrast to externally added Di4 (microinjections in Fig 3B-C, Fig S6D, and Fig S7A were conducted in presence of BSA). Leaflet-specific localization of Di4 after microinjection versus external addition is also supported by their different lifetimes in the PM, as discussed in the results. Also generalized polarization (GP) of Di4, a complementary reporter of membrane packing, showed the same trends as emission lifetime, confirming the robustness of the result (Fig S9).

Finally, we confirmed that Di4 photophysical properties are not affected by charged lipids or transmembrane potential (Fig S8), as previously described[13].

***Materials***: Synthetic lipids were obtained from Avanti. Di-4-ANEPPDHQ (Di4), BSA (fraction V), Phenol red-free MEM, AnxV-Pacific Blue, and dextran-cascade blue (3000 Da) were purchased from Thermo Fisher. NR12S was kindly provided by A. Klymchenko and Mikhail Bogdanov. All other chemicals were purchased from Sigma. Rat basophilic cells (RBL) and NIH 3T3 fibroblasts were purchased from ATCC. Glass micropipettes were from Sutter Instruments and microloaders from Eppendorf. For analysis of red blood cells, 6 channel IBIDI™ slides were used.

### ***Detailed methods: Lipidomics***

*Lipidomics summary:* 300 uL of packed, freshly isolated, intact human erythrocytes from healthy donors with informed consent were treated with 10 I.U. phospholipase A2 (PLA2; from Apis mellifera) or 0.5 I.U. sphingomyelinase (SMase; from Bacillus cereus) in 5 mL isotonic saline solution (50mM Tris HCl, 0.25 mM CaCl2, 0.25mM MgCl2, 150mM NaCl, pH 7.4) for 30 minutes at 30°C to specifically digest only the lipid species present on the exoplasmic leaflet of the PM. After treatment for the indicated time, the cells were fast frozen in liquid nitrogen, and their detailed lipid compositions were analyzed by shotgun electron spray ionization and tandem MS-MS (ESI-MS/MS). All lipidomics were performed at Lipotype GmbH (Dresden, Germany) as described previously [14-19] and detailed below. Lipidomes were prepared from at least 3 independent human donors for all experiments using the following procedures.

*Nomenclature:* The following lipid names and abbreviations are used: ceramide (Cer), cholesterol (Chol), sphingomyelin (SM), diacylglycerol (DAG), lactosyl ceramide (DiHexCer), glucosyl/galactosyl ceramide (HexCer), sterol ester (SE), and triacylglycerol (TAG), as well as phosphatidic acid (PA), phosphatidylcholine (PC), phosphatidylethanolamine (PE), phosphatidylglycerol (PG), and phosphatidylinositol (PI), phosphatidylserine (PS), and their respective lysospecies (lysoPA, lysoPC, lysoPE, lysoPI, and lysoPS) and ether derivatives (PC O-, PE O-, LPC O-, and LPE O-). Lipid species were annotated according to their molecular composition as follows: [lipid class]-[sum of carbon atoms in the FAs]:[sum of double bonds in the FAs];[sum of hydroxyl groups in the long chain base and the FA moiety] (e.g., SM-32:2;1). Where available, the individual FA composition according to the same rule is given in brackets (e.g., 18:1;0-24:2;0).

*Lipid standards*: Synthetic lipid standards were purchased from Sigma-Aldrich (Chol D6), Larodan (Solna, Sweden) Fine Chemicals (DAG and TAG), and Avanti Polar Lipids (all others).

*Lipid extraction for mass spectrometry lipidomics:* Lipids were extracted using a two-step chloroform/methanol procedure [18]. Samples were spiked with internal lipid standard mixture containing: cardiolipin 16:1/15:0/15:0/15:0 (CL), ceramide 18:1;2/17:0 (Cer), diacylglycerol 17:0/17:0 (DAG), hexosylceramide 18:1;2/12:0 (HexCer), lyso-phosphatidate 17:0 (LPA), lyso-phosphatidylcholine 12:0 (LPC), lyso-phosphatidylethanolamine 17:1 (LPE), lyso-phosphatidylglycerol 17:1 (LPG), lyso-phosphatidylinositol 17:1 (LPI), lyso-phosphatidylserine 17:1 (LPS), phosphatidate 17:0/17:0 (PA), phosphatidylcholine 17:0/17:0 (PC), phosphatidylethanolamine 17:0/17:0 (PE), phosphatidylglycerol 17:0/17:0 (PG), phosphatidylinositol 16:0/16:0 (PI), phosphatidylserine 17:0/17:0 (PS), cholesterol ester 20:0 (CE), sphingomyelin 18:1;2/12:0;0 (SM), triacylglycerol 17:0/17:0/17:0 (TAG) and cholesterol D6 (Chol). After extraction, the organic phase was transferred to an infusion plate and dried in a speed vacuum concentrator. 1st step dry extract was re-suspended in 7.5 mM ammonium acetate in chloroform/methanol/propanol (1:2:4, V:V:V) and 2nd step dry extract in 33% ethanol solution of methylamine in chloroform/methanol (0.003:5:1; V:V:V). All liquid handling steps were performed using Hamilton Robotics STARlet robotic platform with the Anti Droplet Control feature for organic solvents pipetting.

*MS data acquisition:* Samples were analyzed by direct infusion on a QExactive mass spectrometer (Thermo Scientific) equipped with a TriVersa NanoMate ion source (Advion Biosciences). Samples were analyzed in both positive and negative ion modes with a resolution of $R_{m/z=200}$=280000 for MS and $R_{m/z=200}$=17500 for MS/MS experiments, in a single acquisition. MS/MS was triggered by an inclusion list encompassing corresponding MS mass ranges scanned in 1 Da increments [16]. Both MS and MS/MS data were combined to monitor CE, DAG and TAG ions as ammonium adducts; PC, PC O-, as acetate adducts; and CL, PA, PE, PE O-, PG, PI and PS as deprotonated anions. MS only was used to monitor LPA, LPE, LPE O-, LPI and LPS as deprotonated anions; Cer, HexCer, SM, LPC and LPC O- as acetate adducts and cholesterol as ammonium adduct of an acetylated derivative [20].

*Lipid identification and quantification:* Data were analyzed with in-house developed lipid identification software based on LipidXplorer (20). Data post-processing and normalization were performed using an in-house developed data management system. Only lipid identifications with a signal-to-noise ratio >5, and a signal intensity 5-fold higher than in corresponding blank samples were considered for further data analysis.

*Lipidomics data processing:* The lipidomic analysis yields a list of >600 individual lipid species and their picomolar abundances. These were processed by first transforming into mol% of all lipids detected. Next, the contaminating TAG and sterol esters were removed from the analysis, and the remaining data was analyzed as mol% of membrane lipids. From here, the datasets were broken down further into class composition. In some cases, the distribution and structural characteristics (e.g. number of carbons or unsaturations in the acyl chains) of the individual species were analyzed. Each class was then compared separately for each individual biological replicate (i.e. the enzymatically treated samples were directly compared to the untreated parallel sample for each individual experiment to control for variance across RBC isolation or human donor).

*Hemolysis measurements*: For all enzyme treatments, hemolysis was monitored to ensure that the RBCs remained intact, and only conditions that yielded no hemolysis were considered for the asymmetry studies (Fig S2). Hemolysis was measured by measuring the absorbance of the supernatant of the enzyme-treated cells at 540 nm on a Tecan plate reader. As an additional test, RBCs were incubated with FITC-dextran (MW=3KDa) during treatment with SMase and imaged on a Nikon A1R laser scanning confocal microscope immediately after.

### ***Detailed methods: Simulations***

*Simulations summary*: Two simulations of symmetric bilayers (i.e. identical compositions in each leaflet) were performed, one with the composition of the inner leaflet and one of the outer leaflet (Supp Table I). For each system, a small patch of membrane 1/9 the area of the final system was built and equilibrated. This small membrane patch was duplicated in a 3x3 array to generate the final system, which was equilibrated further before transferring to Anton2 for production simulation. Initial configurations for the small systems were generated with the CHARMMGUI membrane builder [21], including at least 45 water molecules per lipid and sufficient ions to neutralize the system and set the overall salt concentration to 150 mM, with KCl for the inner leaflet simulation and NaCl for the outer leaflet. The lipids were modeled with the Charmm36 force field [22,23]. All equilibration steps were performed on local resources with NAMD v2.12 [24]. Each small system was minimized by 4,000 steps of conjugate gradient, then heated to 310 K by rescaling velocities to increase the temperature by 3 K every 4 timesteps, followed by another 50,000 timesteps of equilibration at 310 K, rescaling velocities every 100 timesteps. During heating and equilibration the system was coupled semi-isotropically to a Nosé-Hoover/Langevin piston[25,26] with a target pressure of 1.013 bar, a damping timescale of 100 fsec, and period of 200 fsec, and a temperature of 310 K. Long range electrostatics were computed by particle mesh Ewald[27] on a 1.0 Å grid with 4$^{th}$ order interpolation and a tolerance of $10^{-6}$. Long range Lennard-Jones interactions were shifted to zero from 10-12 Å, with both force and potential continuous across the cutoff. The small system was then tiled in a 3x3 array using the periodic unit cell to generate a final system approximately 18 nm x 18 nm, with the final numbers of lipids shown in Supp Table I. These final systems were then equilibrated for an additional 50,000 timesteps by velocity rescaling at 310 K, followed by 20 nsec of Langevin dynamics at 310 K with a damping constant of 0.5 $psec^{-1}$ using a 2.0 fsec timestep, hydrogens constrained by SHAKE with a $10^{-5}$ tolerance.

The systems were then transferred to Anton2 for production with v. 1.37.3c7[28]. The NAMD restart files were converted to dms format using the NAMDtoDMS python script available from the Anton wiki page. Force field information was added with viparr v1.9.1. The systems were integrated with the multigrator with a 2.5 fsec timestep using Martyna-Tobias-Klein for semi-isotropic pressure control at 1.013 bar (no long range dispersion correction) and Nosé-Hoover for temperature control at 310 K[29,30]. Lennard-Jones interactions were cutoff at 9.0 Å. Electrostatics were computed with the k-GSE algorithm[31]. with long-range interactions updated every 3 timesteps. Parameters including cutoffs were chosen by the Anton2 guessers. Each system was run for 5 µsec, with the observables averaged over the last 4 µsec of the simulation.

*Simulation Analysis:* Order parameters. Hydrocarbon chain order was evaluated by computing the 2H NMR order parameter $S_{CD} = |\langle \frac{3}{2}\cos^2\theta - \frac{1}{2} \rangle|$, where $\theta$ is the angle between the carbon-hydrogen bond vector and the membrane normal and the average is taken over both time and lipids of the same type. A single $S_{CD}$ value is computed for every position along both chains for each lipid type, then weighted by the population of each lipid in the mixture to generate the position-specific weighted average $S_{CD}$ curves shown in Fig 2C. The analysis was performed using a modified tcl script kindly provided by Jeff Klauda.

Area per chain. The area associated with each hydrocarbon chain was computed by a Voronoi tessellation of each leaflet, associating a unique area to the center of mass of each chain, then averaging over both chains of the same type and simulation time. The Voronoi analysis was performed with code provided by Alex Sodt[32].

Mean squared displacement. Lipid mobilities were estimated by computing the mean squared displacement (MSD) of each lipid type according to

$MSD(t) = \langle (\boldsymbol{r}(t+t_0) - \boldsymbol{r}(t_0))^2 \rangle$, where $\boldsymbol{r}(t)$ is the two-dimensional position of a single lipid in the membrane plane at time $t$ and the average is taken over both time and lipids of the same type. The MSDs so obtained were then fit to obtain an effective two-dimensional diffusion coefficient. Note that these diffusion coefficients are not expected to be in quantitative agreement with experimental measurements due to finite-size effects in the simulations,[33] but that they can be compared across similarly sized simulations as is done here.

*Hydrocarbon chain packing defects*: The transient exposure of hydrocarbon chains to the aqueous environment was evaluated as described[34]. For each simulation snapshot, the solvent exposure of the hydrocarbon chains was determined using the solvent accessible surface area (SASA) VMD plugin with a probe radius of 3.0 Å. The points representing the SASA of the hydrocarbon chains are then projected onto the membrane midplane separately for each leaflet, generating localized clusters of points representing packing defects. Points are then partitioned into clusters by a breadth first search, connecting all pairs of points that are less than 2.5 Å apart. The area of each defect is then estimated on a grid with a spacing of 0.4 Å by summing the number of occupied pixels for each defect. This procedure is repeated for each simulation snapshot, to generate the histogram in Fig 2E.

### ***Detailed methods: Di4 staining and imaging***

*Cell culture:* Rat basophilic leukemia (RBL) cells were maintained in medium containing 60% MEM, 30% RPMI, 10% FBS, 100 U/ml penicillin, and 100 μg/ml streptomycin. NIH 3T3 cells were maintained in 10% FBS in DMEM with 100 U/ml penicillin, and 100 μg/ml streptomycin.

*Plasmid construction:* All TMD constructs were based on the trLAT backbone previously described [35-37]. The amino acid sequences of WT trLAT is NH2-MEEAILVPCVLGLLLLPILAMLMALCVHCHRLPGS followed by a short linker (GSGS) and monomeric RFP (mRFP). TMD mutants were generated by synthesizing the gene of interest (Genscript) and subsequent cloning of the mutant sequence into the trLAT construct. Mutants were confirmed by sequencing. The TMD sequences used here were:

AexoLcyto: MEEAALAAAALAAALLLLLLLLLLCVHCRLPGSGS

LexoAcyto: MEELLLLLLLLLLLAAALAAAALAACVHCRLPGSGS.

*Transfection and immunofluorescence:* RBLs were transfected by nucleofection (Amaxa) using the protocols provided with the reagents. After 4–6 h of transfection, cells were washed with PBS and then incubated with serum-free medium overnight. To synchronize the cells, 1 h to fixation, the cells were given full-serum medium. Cells were then fixed with 4% formaldehyde for 20 minutes at room temperature. For immunofluorescence labeling of various organelles, the following primary antibodies were used: anti-Rab7 rabbit monoclonal antibody (late endosome; Cell Signaling Technology), anti-Rab5 rabbit monoclonal antibody (late endosome; Cell Signaling Technology), anti-Giantin rabbit polyclonal antibody (Golgi; Abcam), and anti-LAMP1 rabbit polyclonal antibody (lysosome; Abcam). Fluorescent (Atto 488) secondaries were purchased from Invitrogen.

*PM localization:* PM localization was quantified by an automated imaging protocol[35]. Briefly, transfected RBL cells were surface-labeled by a membrane-impermeable biotinylation reagent (Biotin-NHSLC) at 1 mg/ml in PBS for 20 min at 4 °C. Cells were fixed with 4% paraformaldehyde, and the PM were fluorescently labeled by streptavidin-Alexa488 (10 μg/ml for 20 min). Cells were imaged on a Nikon A1R confocal microscope using a ×60 Apochromat oil immersion objective. PM localization was quantified by a custom image processing protocol (MATLAB) in which the fluorescence intensity of the protein construct that was co-localized with the plasma membrane stain was divided by the total fluorescence intensity in the whole cell.

*Leaflet-selective staining of plasma membrane leaflets*: Outer leaflet staining of cultured cells was achieved by incubating cells at 4°C for 8 min with Di4 at 1 μg/ml in staining buffer (10 mM HEPES, pH=7.4, 150 mM NaCl, 2 mM $CaCl_2$). Cells were washed twice in the same buffer at ambient temperature before imaging in phenol red-free MEM. For dextran chase assays, cells were stained in the same manner.

To specifically stain the inner leaflet, Di4 was microinjected directly into the cytoplasm. Briefly, cells were kept in MEM media (no phenol red) containing 1.5% BSA to capture any dye leakage from the microinjection needle, which prevents staining the outer leaflet of surrounding cells (compare Fig S5C-D and Fig S6) and also extracts any external PM dye (Fig S5B). The microinjection solution was prepared by evaporating 4 μg of Di4 (dissolved in EtOH) under nitrogen flow, re-dissolving in 8 μl microinjection buffer containing 10 mM potassium phosphate (mono and dibasic to adjust at pH = 7) and 100 mM KCl. We then centrifuge at 10,000 xg for 10 min to remove undissolved material.

Microinjection needles were pulled from glass micropipettes (1 mm outer, 0.56 mm inner diameter) and loaded by micro-loaders (2 µl). Microinjection was performed using a Transjector 5246 (Eppendorf) installed on a light microscope to visualize the cells. After microinjection, cells were bathed in MEM medium (no phenol red) without BSA and imaged as below. Some microinjected cells were further stained from the outside after imaging (outside/inside staining) by adding Di4 to the bathing medium. To exclude apoptosis or membrane disruption by microinjection, cells were stained with Annexin V (AnxV-pacific blue) after FLIM and/or GP imaging; all AnxV-positive cells were excluded from the analysis.

To remove outer leaflet Di4 from labeled cells, cells were incubated 3 times for 10 min with 1.5% BSA in MEM media at 4°C (to prevent endo- or exocytosis). To test whether BSA itself affects membrane properties, RBLs were incubated 3x for 10 min with 1.5% BSA at 4°C, washed with PBS, subsequently stained with 1 µg/mL Di4 at 4°C for 8 min, and imaged via FLIM (as described below).

Isolation and treatment of red blood cells for imaging: RBCs were freshly isolated from human donors and diluted at $5e^6$ cells/ml in Ringer's solution. Cells were then stained with Di4 (1 ug/ml) at 4°C for 8 min or NR12S (100 nM) for 10 min at at 37°C, centrifuged at 2000g for 5 min, resuspended in Ringer's solution. IBIDI™ slides were used for FLIM and GP imaging. To induce membrane scrambling, the RBCs were treated with 10 µM PMA for 10 min under constant mixing at 300 rpm. RBCs were stained with AnxV-pacific blue, only after imaging Di4 imaging, to verify scrambling and PS exposure while not interfering with Di4 spectra/lifetime (e.g. via FRET).

Confocal spectral imaging and lifetime imaging microscopy (FLIM): Confocal and lifetime microscopy were performed on a Nikon A1 laser scanning microscope with an integrated Picoquant time-correlated single photon counting (TC-SPC) system (Berlin, Germany). Generalized polarization (GP)-imaging was done as previously described [14,15]. Briefly, emission was collected at I1=595 and I2=700 nm and GP was calculated as $GP=(I_1-G*I_2)/(I_1+G*I_2)$ where G, the instrumental response factor, was determined according to protocol[9].

For FLIM, Di4 emission was collected at >560 nm and the instrument response function (IRF) was determined with a saturated erythrosine B and KI solution at pH=10 according to the manufacturer's (Picoquant) protocol. Di4 images were acquired using 20 MHz pulse frequency. The photon count rate was kept under 10% of the pulse rate by adjusting a manual shutter, and enough frames were acquired to obtain at least $10^4$ photons cumulative signal intensity. The fluorescence decay curves were fitted to a bi-exponential re-convolution function adjusted to the IRF and the average lifetime was calculated and represented in the FLIM images as $\tau_{Di4}$.

Dextran chase experiment: RBL and 3T3 cells were incubated with dextran-cascade blue (0.5 mg/ml) for 2 h at 37°C. Cells were then washed twice with HEPES buffer (10 mM HEPES, 150 mM NaCl, 2 mM $CaCl_2$, pH 7) and stained on the inner or outer plasma membrane leaflet by Di4 (see selective staining). After staining, cells were incubated for 15, 30, or 60 min (chase) at 37°C in MEM media, allowing dextran to accumulate in endosomes and Di4 to access endosomal membranes.

Preparation of GUV: GUV of DOPC/DPPC/Chol were prepared by electroformation as described previously [38]. Briefly, 1 µl of the lipid solution (1:2 methanol/chloroform at 2.5 mg/ml) was applied to Pt electrodes of an electroformation chamber. The solution was dried in vacuum for 3 h, then rehydrated in a 0.4 M sucrose. Electroformation was performed at 500Hz and 2.5 V for 2 h at 50°C. GUVs containing charged lipids (DOPC/DOPS, 4:1) were also prepared by hydration swelling to avoid potential issues with electroformation[39,40]. Briefly, glass slides were incubated with 1% agarose ultra-low gelling temperature (Sigma-Aldrich) for 2 min at ambient temperature. Slides were then heated at 40°C. After drying, 30 µl of lipid solution (4 mg/ml in CHCl3/MeOH 9:1) was spread over the film and evaporated. The slide was kept in vacuum overnight and rehydrated with 0.4 M sucrose solution the next day at ambient temperature for 2h. All GUVs were later stained by adding Di4 at 250 nM final concentration and incubated for 5 min. 10 µl of stained GUVs were rediluted into 200 µl of 0.4 M glucose solution (containing 0.5% agarose to prevent moving of GUV during FLIM) and imaged. To show efficient integration of DOPS into GUVs, we stained GUVs with Anx-V-640 after formation and visualized the fluorescent GUV in a fluorescent microscope (data not shown). To test the lifetime of Di4 under various buffer conditions, the GUVs were diluted into the following solutions (all brought to 400mOsm/L with glucose) prior to imaging: intracellular buffer (20 mM HEPES, 110 mM KCl, 10 mM NaCl, 2 mM MgCl2, 5 mM KH2PO4, pH 7.3), extracellular buffer (20 mM HEPES, 5 mM KCl, 140 mM NaCl, 1 mM MgCl2, 1 mM CaCl2, 5 mM KH2PO4, pH 7.3), 20 mM HEPES pH 5.2, and 20 mM HEPES pH 7.0.

Determination of Di4 flipping rate: One milliliter of a chloroform/methanol (2:1) solution containing 10 mM DOPC +/- Di4 (molar ratio 1:200) was evaporated in a rotary evaporator under vacuum for 1 h and stored in a vacuum dessicator overnight. Multilamellar vesicles (MLV) were made according to the freeze-thawing method by hydrating the lipid film with buffer containing 10 mM Tris, 400 mM NaCl, pH 7.4. The MLV were then extruded 15 times through a 200 nm filter to obtain LUVs with a uniform size distribution. Size distribution was verified by dynamic light scattering and phospholipid content was determined by the inorganic phosphate assay [41]. To determine the flipping speed, LUV were diluted to 100 µM phospholipids and we measured the magnitude of Di4 quenching ($\lambda_{ex}$= 480, $\lambda_{em}$=650 nm) by potassium iodide (KI; 800 mM) over time at 20°C. To that end, LUVs were post-labeled with Di4 by incubating pre-made LUVs with 500 nM Di4 for 8 min incubation at 4°C. KI was then added at different time points after labeling and the resulting fluorescence drop was measured in a fluorimeter (Clariostar, BMG Labtech, Ortenberg, Germany). LUVs prepared with Di4 on both leaflets (i.e. Di4 included during vesicle production) were used as a control to estimate the amount of quenching when equal abundance of Di4 was present on both leaflets. When these vesicles were sonicated in the presence of KI, we observed similar quenching as for outside-labeled vesicles.

Quenching efficiency was calculated as:

$QE = 1 - F_{quenched}/F_{unquenched}$

where $F_{quenched}/F_{unuenched}$ is the ratio of fluorescence intensity of Di4 in presence and absence of KI, respectively. The percentage of Di4 on the outer leaflet at various timepoints was calculated as:

$\%Di4_{outside} = 100 * (QE_{Di4,outside} / 2* QE_{Di4,both})$

where $QE_{Di4out}$ is the quenching efficiency of Di4 in LUVs where Di4 was only added to the outer leaflet and $QE_{Di4, both}$ is the quenching efficiency of LUVs in which Di4 was present at both leaflet. The factor 2 comes from the fact that only half of the Di4 molecules are on the outer leaflet in that configuration. Three independent experiments were done with triplicate measurements for quenched and unquenched LUVs.

*Fluorescent Correlation Spectroscopy:* We measured the diffusion coefficient of GPI-GFP and SH4-mNeonGreen on CHO cells using fluorescence correlation spectroscopy. CHO cells were seeded on glass slides (#1.5) two days before the measurement. One day before the measurement, cells were transfected using Lipofectamine 3000 using the protocol supplied by the manufacturer. Both proteins were localized almost exclusively on the PM (not shown). FCS on transfected cells was carried out at room temperature using Zeiss LSM 880 microscope, 40X water immersion objective (numerical aperture 1.2). The laser spot was focused on the basal membrane of the cells by finding the focal plane of maximum fluorescence intensity. Then, 3-5 curves were obtained for each spot (five seconds each). The obtained curves were fit using the freely available FoCuS-point software using 2D and triplet model.

*Bioinformatics*: We analyzed all transmembrane domains from single-pass transmembrane proteins annotated in the Uniprot database as previously described [37]. Lipid accessible surface area (ASA) of TMDs was calculated from structural / statistical predictions based on amino acid sequence, as described previously [42]. The exoplasmic/luminal and cytoplasmic portions of the TMD were defined as first and second half of the transmembrane domain according to the annotated orientation of the protein in the plasma membrane (i.e. N-terminal or C-terminal outside). For TMDs with an uneven number of amino acids, the ASA of the amino acid in the center of the membrane was divided by 2 and half of its ASA was attributed to each half of the exo/cytoplasmic side.

**Supplementary Figures**

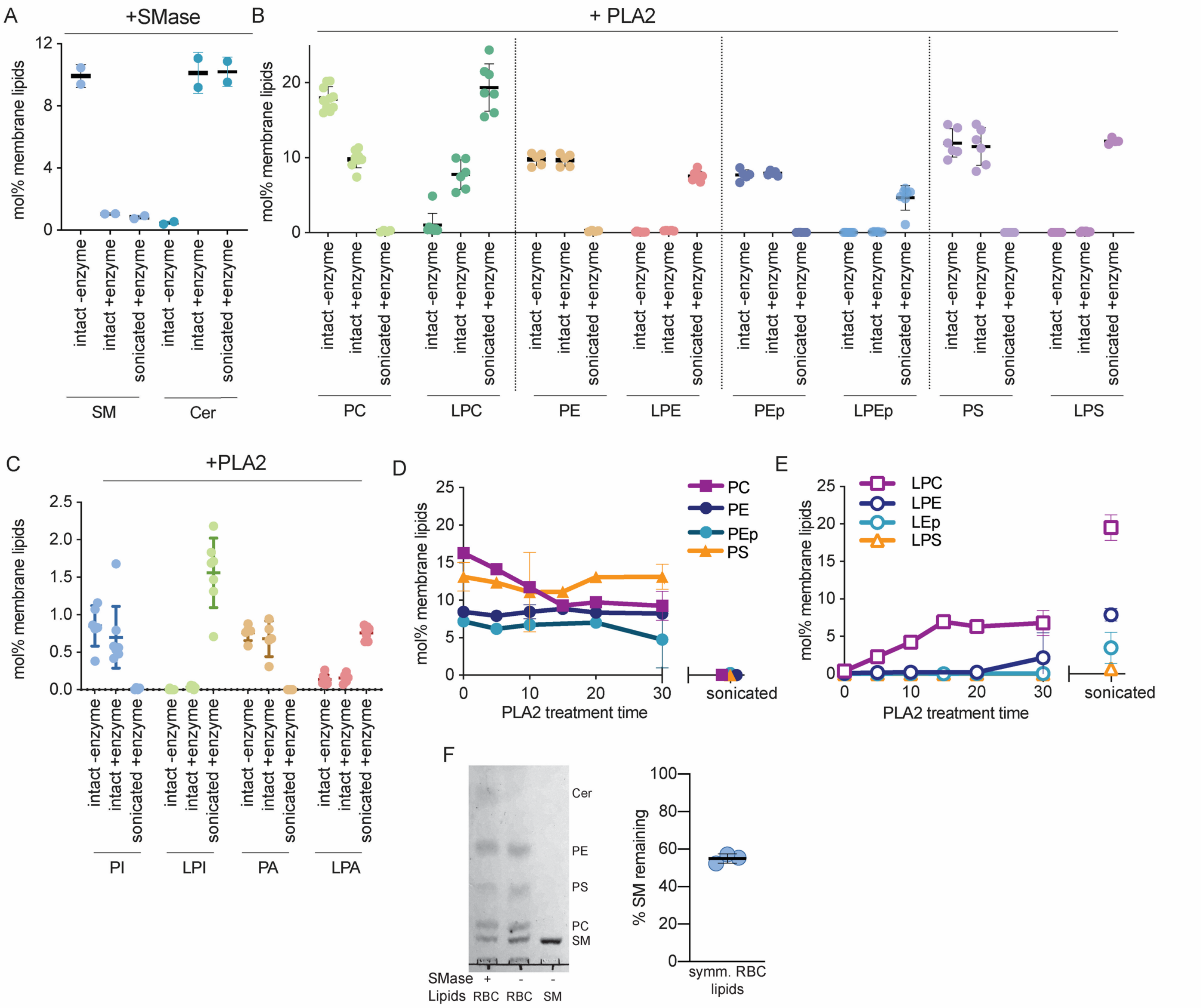


**Fig S1 – Lipidomics of enzyme treated erythrocytes (RBCs).** (A) SMase treatment of intact RBCs degraded ~90% of SM. Treatment of sonicated cells degraded essentially all SM, confirming enzyme potency. Notably, Cer (the product of the SMase reaction) was produced in quantitative agreement with the consumption of SM. Because SM and Cer are quantified using distinct internal standards, the excellent agreement between product generation and reactant consumption confirms quantitative accuracy of the lipidomic measurements. (B-C) Lipidomics after PLA2 treatment of intact and sonicated RBCs. PLA2 did not degrade measurable amounts of PS, PE, PE O-, PI, or PA in intact cells, but completely degraded those in cells disrupted by sonication. These results suggest that such lipids are exclusively on the inner PM leaflet. Again, the quantitative agreement between phospholipids consumed by the reaction and lysolipids generated supports the accuracy of the measurements. (D) Lipid hydrolysis by PLA2 time course. Progressive degradation of PC produces progressive generation of (E) LPC. The reaction saturates at 15 minutes, suggesting all accessible phospholipids are degraded. All other enzyme treatments were done for 30 min. (F) Lipids were extracted from RBCs and reconstituted into 100 nm liposomes. These liposomes were then treated with SMase and the results analyzed via thin layer chromatography (SM lipid standard in rightmost lane). The intensity of the SM band was reduced by enzymatic treatment (left lane). This effect was quantified by densitometry, normalizing to PC, PE, and PS in the same sample. SM was reduced by ~50% by treatment of the symmetric liposomes with SMase, as would be expected from near-complete digestion of all outer leaflet SM. Each data point represents results from a single RBC isolation and treatment; 3 different donors in total were used, some for multiple isolations. As there were no notable differences between donors or isolation days, all samples were combined.

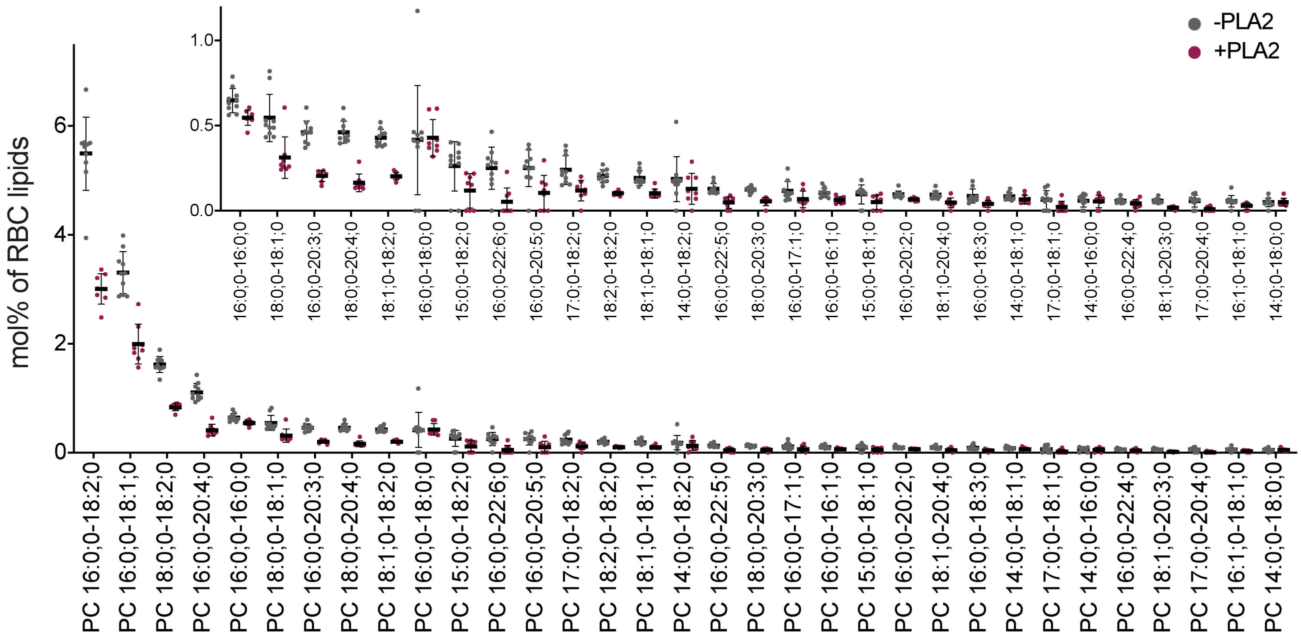


**Fig S2 – Variance in the lipidomic data**. Shown are the individual data points and average ± SD of n>8 individual MS analyses from n>3 human donors. All species of PC above 0.05 mol% are shown for both untreated and PLA2-treated samples. Inset shows the more minor species with an expanded y-axis range.

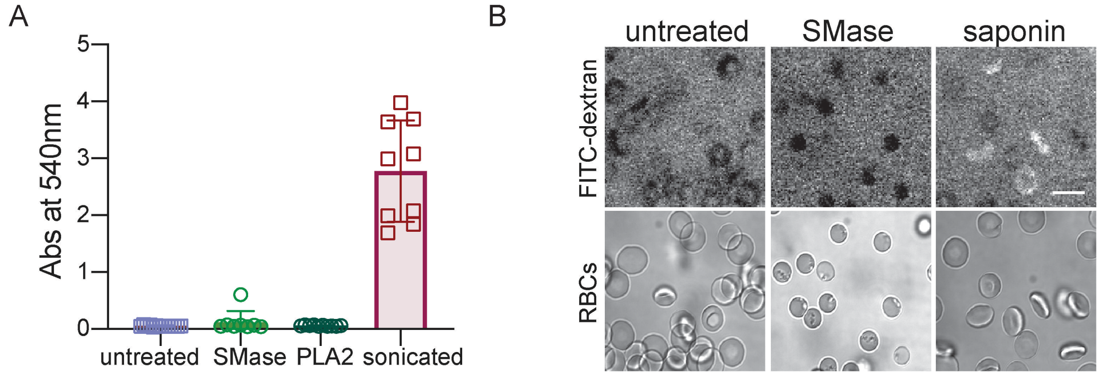


**Fig S3 – Lack of RBC hemolysis following SMase or PLA2 treatments**. (A) Absorbance of supernatant at 540 nm after pelleting RBCs. Neither SMase nor PLA2 treatments lead to any observable absorbance above untreated RBCs, suggesting no leakage of hemoglobin, thus intact erythrocytes. Leakage of hemoglobin following sonication serves as the positive control. (B) Both untreated and SMase-treated RBCs remain impermeable to FITC-dextran (visible by bright background fluorescence and dark cell interior), while saponin-permeabilized cells are completely penetrated, as evidenced by approximately equal FITC-dextran signal inside the cells as outside. These results illustrate that SMase-treated RBCs remain intact and impermeable. Top row shows FITC-dextran fluorescence; bottom row shows pseudo-DIC image to visualize cells. Scale bar is 10 µm.

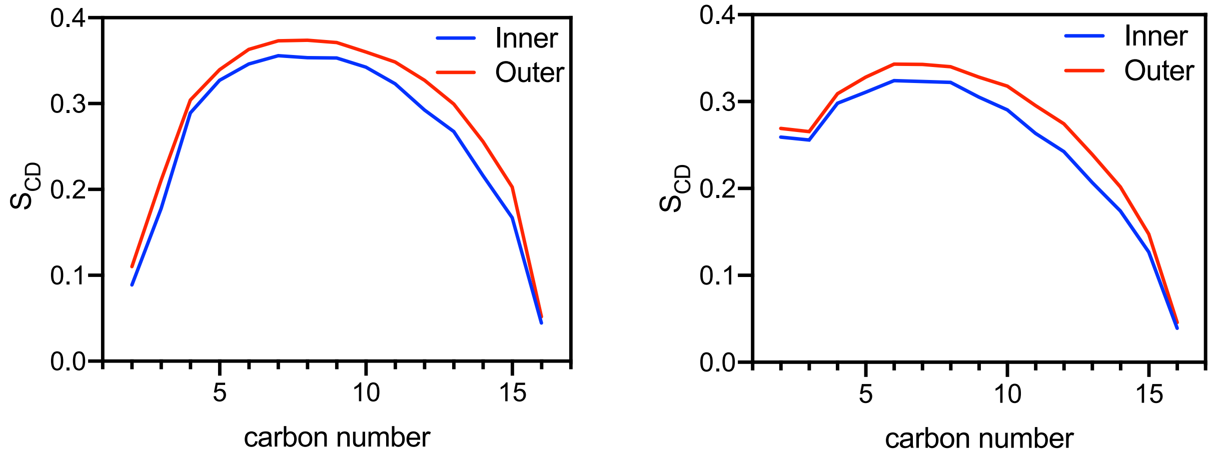


**Fig S4 – Computed deuterium order parameters ($S_{CD}$) of lipids in inner versus outer PM simulations.** (A) the fatty acyl chain of palmitoyl sphingomyelin (PSM). (B) the sn-1 chain of 1-palmitoyl-2-linoleoyl PC (16:0/18:2 PC).

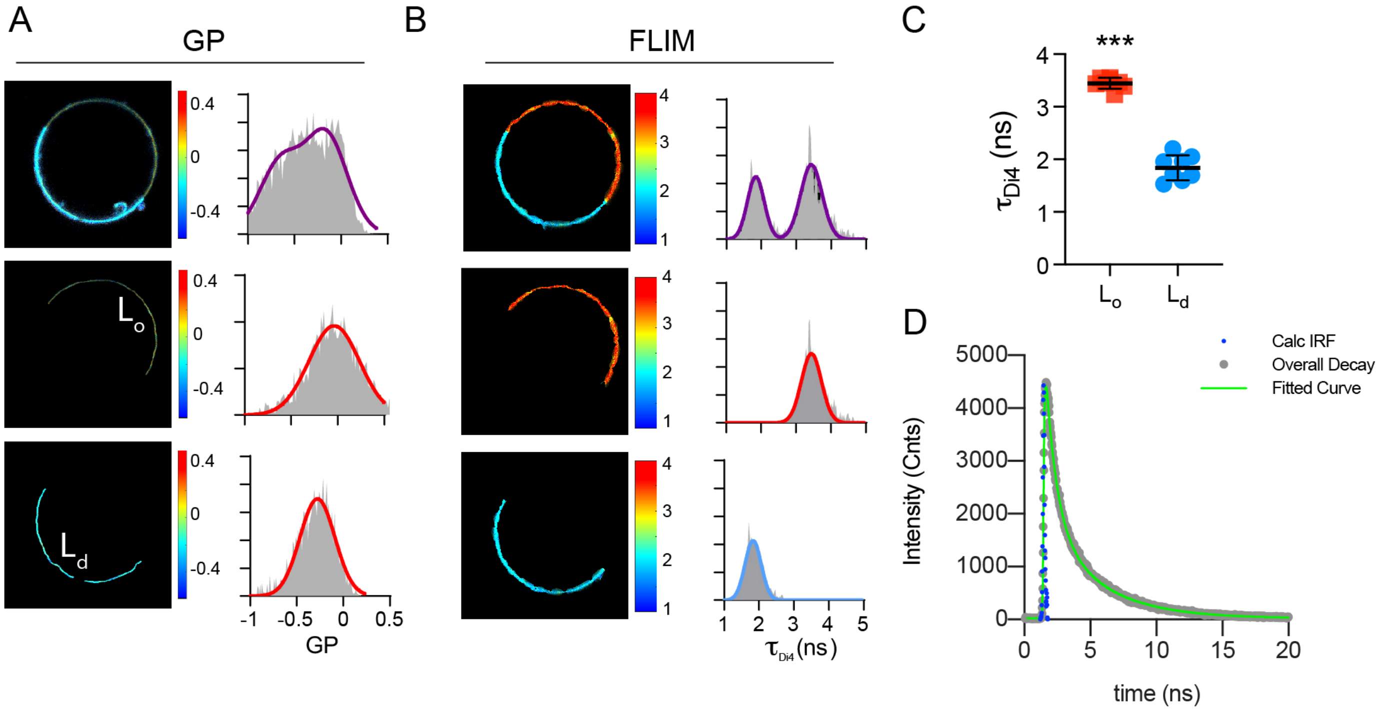


**Fig S5 – Characterization of Di4 for measuring lipid packing differences.** (A) Average GP and lifetime images of Di4 in a phase separated GUV composed of DOPC/DPPC/Cholesterol (40:40:20). The intensity-weighted average lifetime and GP histograms were fit with either a double (whole GUV, GPMV violet line) or single ($L_o$ or $L_d$ phase, red and blue line) Gaussian, respectively. (C) Average lifetime of Di4 in $L_o$ and $L_d$ in GUVs. (D) Representative lifetime curve of Di4 from outer leaflet-stained RBL cells showing the instrument response function (IRF), overall decay data points, and the two-component regression fit ($\tau_{Di4} = 3.15$ ns).

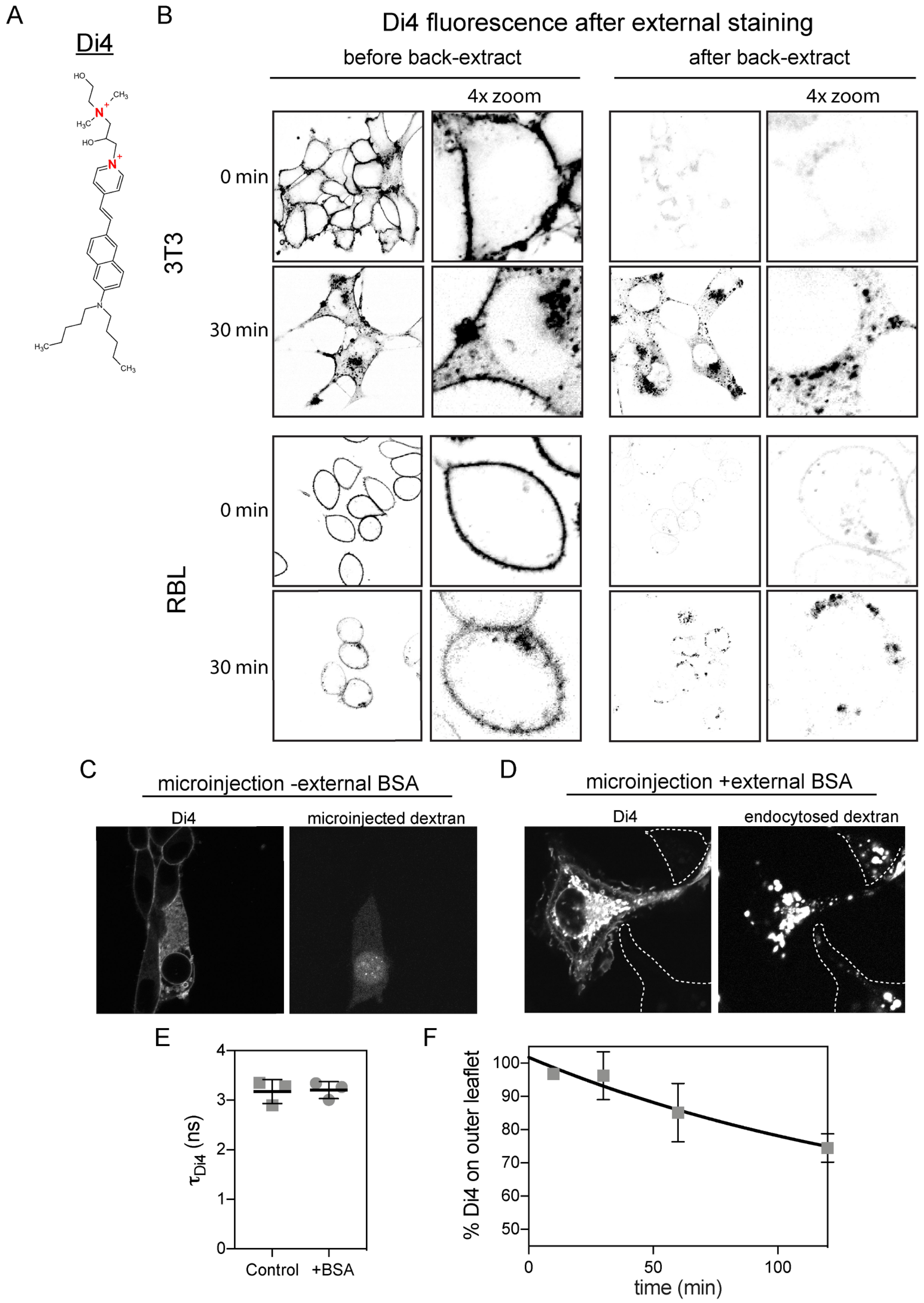


**Fig S6 – Characterization of leaflet selective PM staining by Di4.** (A) di-4ANEPPDHQ (Di4) chemical structure containing 2 positively charged quarternary ammonium groups (red). (B, left) External staining of cells at the PM after staining with Di4. Time 0 is imaging immediately after Di4 incubation and wash. The bottom panels in each group show cells after 30 min incubation at 37°C to allow dye internalization into endosomes. (B, right) Addition of BSA leads to near complete extraction of Di4 suggesting it is solely in the exoplasmic PM leaflet at time 0. After dye is endocytosed (30 min incubation at 37°C), it is not extractable by BSA; however, the dye remaining on the PM is completely extracted. This observation suggests minimal flipping across the bilayer on our experimental time scale. (C) Microinjection of Di4 in the absence of external BSA leads to staining of both the microinjected cell (visualized by co-microinjected dextran-cascade blue at 4 mg/ml) and surrounding cells due to leakage of dye from micropipette. (D) Inclusion of BSA into the bathing media eliminates off-target staining (other cells in the field visualized by endocytosed fluorescent dextran). There remains robust staining of the PM in the target cell because the dye is on the inner leaflet which is not accessible to BSA back-extraction. The same effect (i.e. no extraction of internal membranes by BSA after microinjection) was observed after incubating the cells for up to 60 min at 37°C. (E) BSA treatment itself had no effect on the properties of the PMs. For this experiment, RBLs were incubated with 1.5% BSA at 4°C (3 times for 10 mins), washed with PBS, the *subsequently* stained with Di4 as described in the Materials and Methods. The lifetime of Di4 on the PM was not affected by 'pre-treatment' with BSA, suggesting that BSA itself does not have an effect on membrane properties. (F) To determine the intrinsic flipping rate of Di4 under controlled conditions, we constructed symmetric liposomes (large unilamellar vesicles; LUVs) containing Di4. We then incubated the LUVs with potassium iodide (KI) to quench the fluorescence of externally accessible Di4 (see Methods). Probe flipping results in reduction of quenching efficiency, because KI cannot cross the membrane. Although some flipping was observed, the magnitude was minimal through the time course of the experiment. Fitting to first-order kinetics suggested a flipping rate of ($k_{flip}$=0.0056 min$^{-1}$, C.I.$_{(95\%)}$ = 0.0039-0.0075 min$^{-1}$) with resulting time constant for loss of asymmetry being ~164 min. Thus, on the maximal time-scale of our experiments (60 min), ~14% of the probe would be expected to flip between leaflets, which would only slightly influence the measurements of overall leaflet order. Moreover, we point out that these experiments were done in cholesterol-free liposomes, while the cholesterol-rich PM would be expected to be even less amenable to flip-flop.

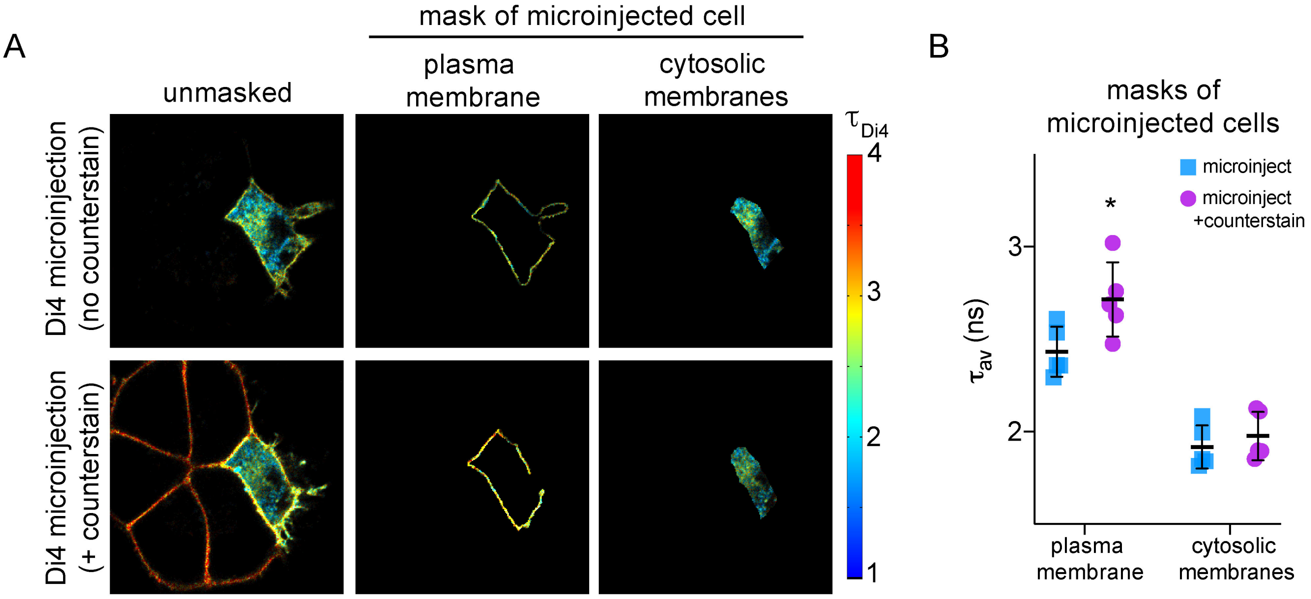


**Fig S7 – Leaflet selective lifetime of Di4.** (A, top row) A single RBL cell was microinjected with Di4 in the presence of external BSA and imaged via FLIM. Without BSA in the medium, surrounding cells are always stained by Di4 leaking out of the microinjection pipette (e.g. see Fig S6C). However, inclusion of BSA leads to Di4 extraction from the outer leaflet, such that only the microinjected cell is visible. (A, bottom row) After imaging and washing away the external BSA, the same cells were re-incubated with Di4 and imaged. Shown are Di4 FLIM images, with PM (middle) and cytosolic membrane (right) masks of the microinjected cell. In the bottom row (after external labeling), the microinjected cell has Di4 in both PM leaflets, whereas the surrounding cells are only stained on the external leaflet, as revealed by the distinct Di4 lifetimes in the PMs. (B) Di4 lifetime of the PM and cytoplasmic membranes of microinjected cells before and after external staining with Di4. Di4 lifetime on the PM is increased by external staining, due to signal coming from both the internal and external PM leaflets. On the other hand, the lifetime of the internal membranes is unaffected.

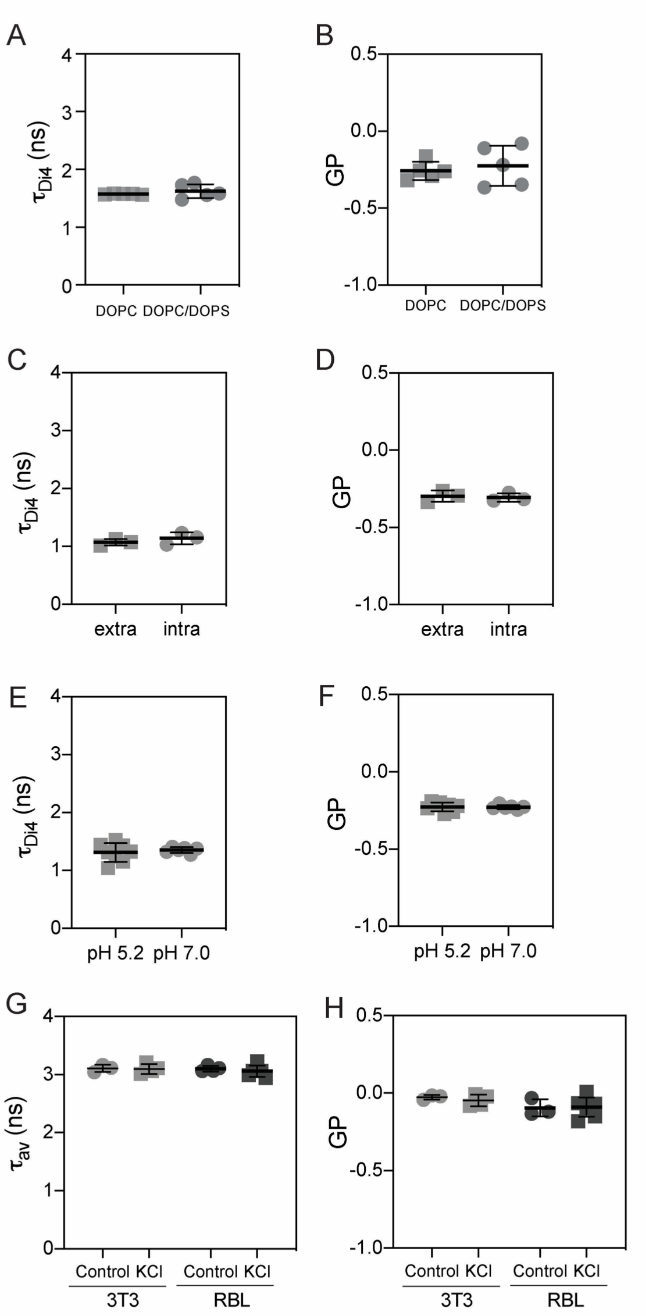


**Fig S8 – Control experiments to exclude the possibility that Di4 is sensing differences in leaflet charge density, ion composition, pH, or membrane potential**. (A) FLIM and (B) GP imaging of Di4 in GUVs composed of either pure DOPC or DOPC with 20% charged lipid (DOPC/DOPS ; 4:1). (C) FLIM and (D) GP of Di4 in GUVs composed of DOPC and incubated in extra- or intracellular buffer. (E) FLIM and (F) GP of Di4 in GUVs composed of DOPC and incubated in 20mM HEPES, pH 5.2 or pH 7.0. Neither lifetime nor GP was affected by the negatively charged DOPS, pH, or ionic strength/composition of buffer suggesting no effect of charged lipids or relevant environmental conditions on Di4 photophysical properties. (G) FLIM and (H) GP imaging of Di4 stained cells before and after membrane depolarization with 100 mM KCl. We observed no effect of membrane depolarization on either parameter, confirming previous observations that surface charge and membrane potential do not significantly affect Di4 lifetime and emission spectra [13]. Finally, Di4 fluorescent properties are insensitive to protein content of the membrane [43], excluding asymmetric protein density as the explanation for our observations.

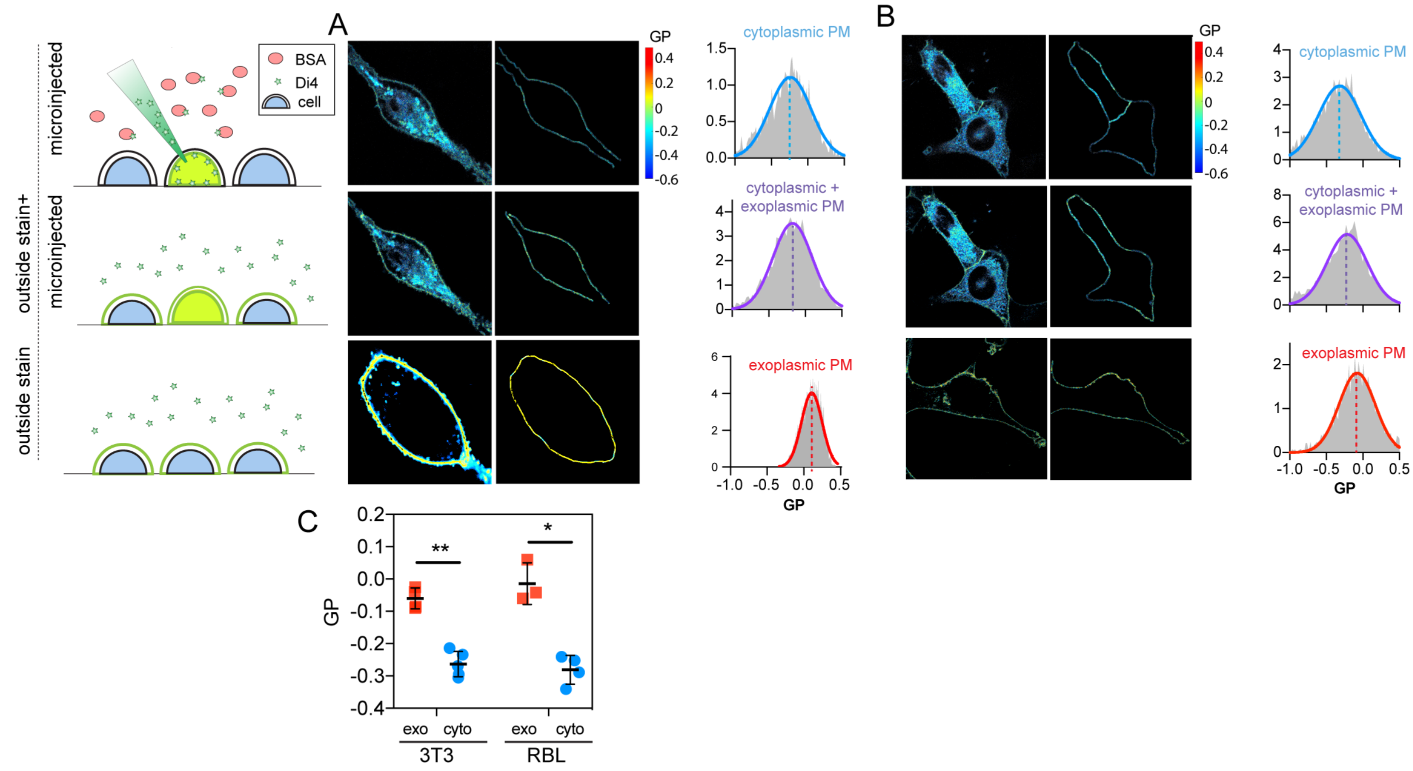


**Fig S9 – Di4 GP imaging confirms the order differences between inner and outer plasma membrane leaflets.** Exemplary GP images of (A) mast cells or (B) 3T3 fibroblasts showing (left) whole cells, (middle) PM masks and, (right) intensity-weighted histograms of the PM mask in cytosolic leaflets (microinjected), both leaflets (outside stain+microinjected), and outer leaflet (outside stain). (C) Average GP of plasma membrane masks of microinjected (blue) vs. outer leaflet stain (red) in both cell types.

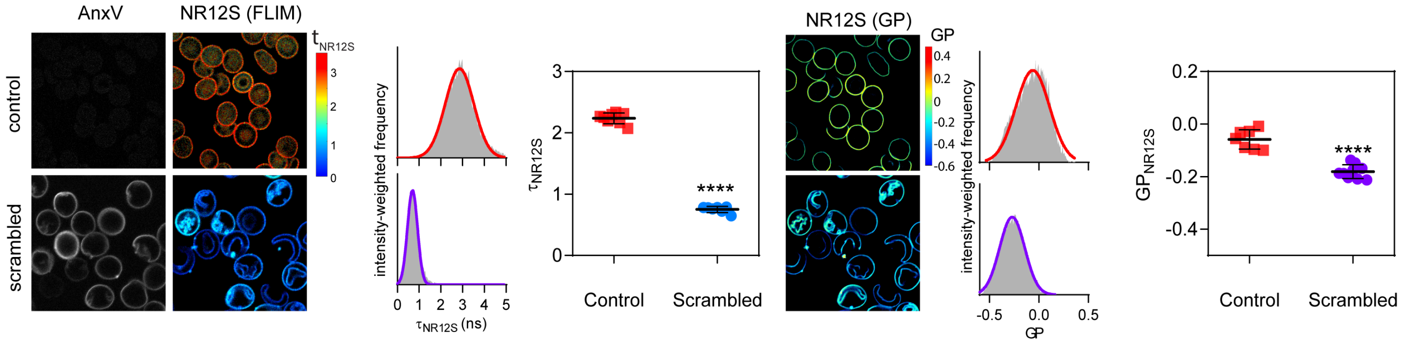


**Fig S10 – Confirmation of reduction of lipid packing in scrambled vs. unscrambled RBCs probed by leaflet selective, order sensitive dye NR12S.** AnxV staining, FLIM and GP imaging of NR12S in untreated (control) vs. PMA 10 µM (scrambled) RBCs. Treatment with PMA induces membrane scrambling, as evidenced by wholesale PS exposure. This scrambling leads to dramatic reduction of NR12S lifetime and GP, suggesting that the outer leaflet of the RBC PM becomes much less packed. These observations suggest that the inner leaflet is much less packed than the outer in resting RBCs. Importantly, AnxV was added right after FLIM or GP-imaging of Di4 to confirm the loss of asymmetry in the imaged cells but avoid any interference with Di4 or NR12S emission.

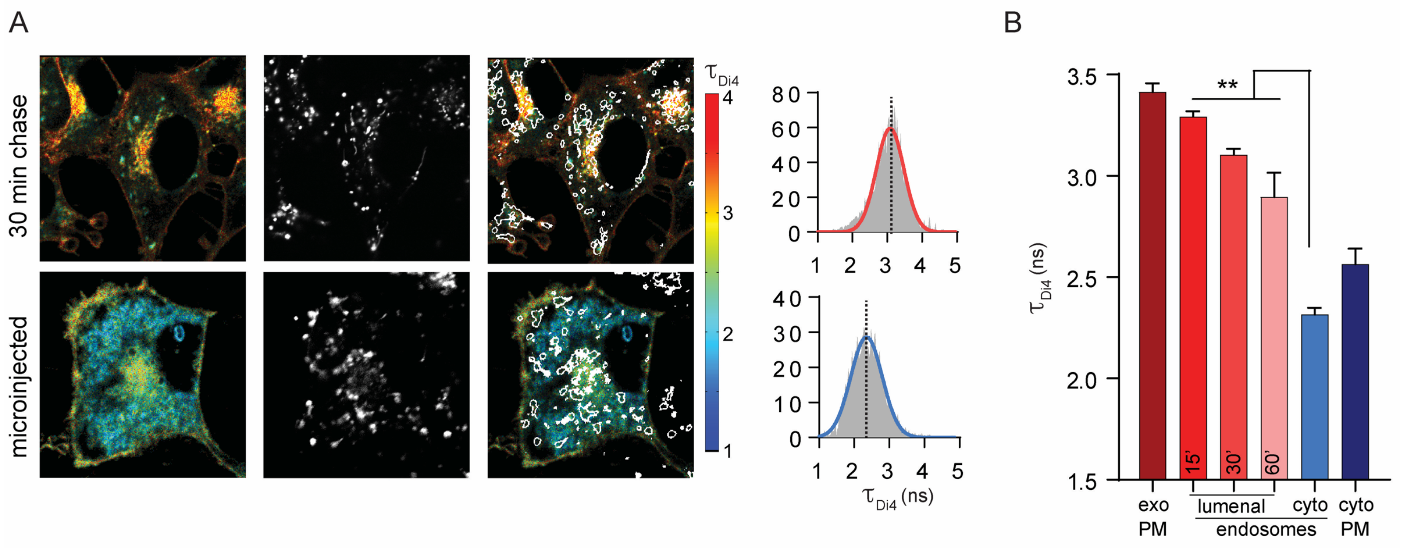


**Fig S11 – Endosome asymmetry in 3T3 fibroblasts**. (A) Exemplary FLIM images of Di4 lifetime and dextran fluorescence following a two-hour incubation with fluorescent dextran, 10-minute staining with Di4, then a 30-minute incubation without either label to "chase" dyes into endosomes. The localization of dextran (middle) is used to create an endosomal membrane mask (right images) to derive intensity-weighted histograms of $\tau_{Di4}$ (right). (B) The high lifetime (i.e. lipid packing) of the exoplasmic PM leaflet is maintained up to 60 min after endocytosis.

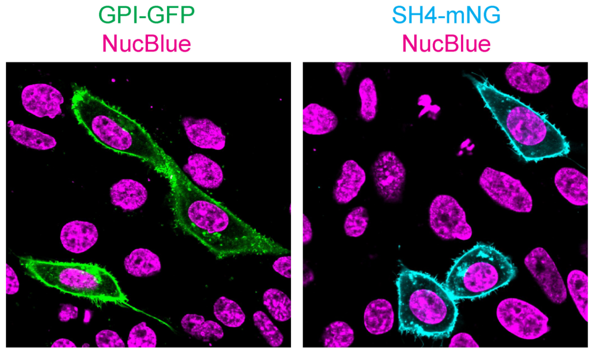


**Fig S12 – GPI-GFP and SH4-mNG localize to the plasma membrane.**

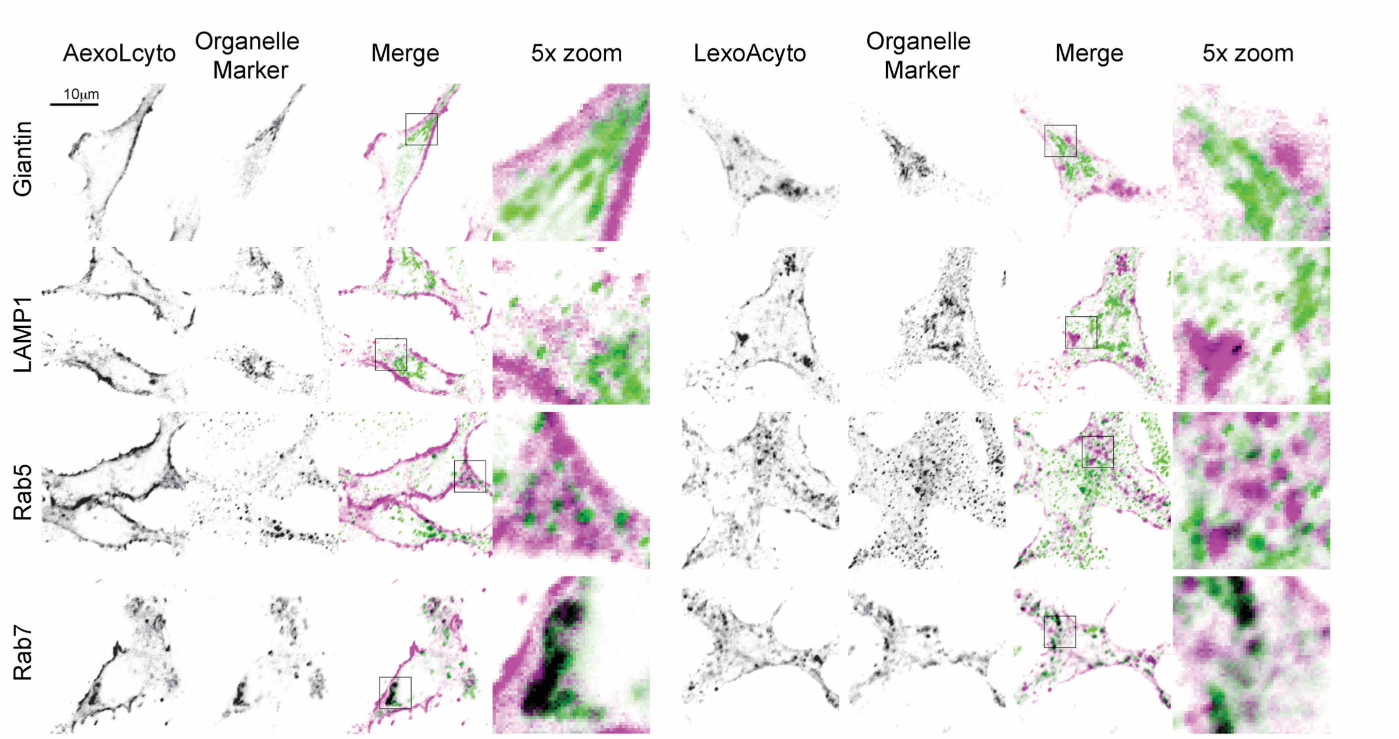


**Fig S13 – Subcellular localization of AexoLcyto-RFP and LexoAcyto-RFP.** RBL cells were transfected with the indicated plasmid, fixed, and immunofluorescence performed. Images were acquired at 100x on a Nikon A1R confocal microscope.

**Supplementary text:**

Disparities in biophysical properties between the two leaflets of the PM in living cells have long been proposed [44], but remain ambiguous due to experimental limitations and inconsistencies. For example, EPR studies found that spin-labeled phosphatidylcholine (PC) was found in a more rigid environment than amine-headgroup lipids, implying that the inner leaflet of the PM is more fluid [18,1945,46]. In opposition, several studies with various fluorescent probes reported a more fluid outer leaflet [47-51]. Similar inconsistencies have been reported by measurements of diffusion with NBD-labeled lipids, with some inferring slower diffusion of outer leaflet components [52,53], while others reported faster diffusion in the outer leaflet at low temperatures and minimal differences at physiological temperature [54]. Similarly, NBD-labeled phosphoinositides diffused faster in the outer leaflet compared to the inner [55].

Despite these significant investigations, the presence and nature of biophysical asymmetry in mammalian membranes has remained unresolved because of several limitations inherent to methodologies that rely on probes based on biological lipids and/or fluorescence quenchers: (1) fluorescence quenchers are toxic and potentially membrane disruptive. Further, their membrane impermeability is questionable and is rarely tested. (2) The localization of the probe in the cell is often not directly known or reported; PM localization has been often assumed but not usually directly imaged. (3) Limited direct knowledge of probe asymmetry. (4) Possible interactions between probes and other lipids, proteins and the cytoskeleton. (5) Interconversion or degradation of fluorescent lipid analogs [56]. In contrast, the probes used here (Di4 and NR12S) are intended to be biologically inert, in that they are not expected to be metabolized like native lipids or to be toxic or disruptive. Direct imaging confirms that they remain on the PM, where their flipping rate can be directly quantified.

### Supplementary Tables

**Supplementary Table I – Compositions of INNER and OUTER PM simulations**.

**(A) OUTER**

| lipid | structure | number |
|---|---|---|
| PSM | SM 34:1;2 | 81 |
| PLPC | PC 16:0;0-18:2;0 | 99 |
| NSM | SM 42:2;2 | 63 |
| LSM | SM 42:1;2 | 54 |
| SOPC | PC 18:0-18:1 | 45 |
| PAPC | PC 16:0-20:4 | 36 |
| SAPS | PS 18:0-20:4 | 9 |
| PLAS | PE-O 18:1-20:4 | 18 |
| CHOL | cholesterol | 270 |

**(B) INNER**

| lipid | structure | number |
|---|---|---|
| PAPS | PS 16:0-20:4 | 90 |
| PLPC | PC 16:0-18:2 | 63 |
| POPC | PC 16:0-18:1 | 27 |
| PLQS | PE-O 18:1-22:4 | 63 |
| POPE | PE 16:0-18:1 | 18 |
| PDPE | PE 16:0-22:6 | 54 |
| OAPE | PE 18:1-20:4 | 27 |
| PLAO | PE-O 18:2-20:4 | 18 |
| DPPC | PC 16:0-16:0 | 18 |
| PSM | SM 34:1;2 | 9 |
| OAPS | PS 18:0-20:4 | 9 |
| PIP2 | PIP2 18:1-20:4 | 18 |
| CHOL | cholesterol | 279 |

**Supplementary Table II – Compiled inner and outer leaflet lipidomes of RBCs**

**(A) Outer leaflet compiled lipidome**

| Category | Notes | representative lipid | mol% of outside phospholipids | scaled to 100% |
|---|---|---|---|---|
| 1 | a,b | PC 16:0-18:2 | 19.7 | 22.2 |
| 2 | a | SM 34:1;2 | 19.8 | 22.4 |
| 3 | | SM 42:2;2 | 16.4 | 18.5 |
| 4 | c | SM 42:1;2 | 12.8 | 14.4 |
| 6 | e | PC 16:0-20:4 | 8.6 | 9.7 |
| 5 | | PC 18:0-18:1 | 7.5 | 8.4 |
| 9 | d | PC 18:1-18:2 | 1.8 | 2.0 |
| 7 | e | PS 18:0-20:4 | 1.3 | 1.5 |
| 8 | e | PI 18:0-22:4 | 0.8 | 0.9 |

**(B) Inner leaflet compiled lipidome**

| category | notes | representative lipid | mol% of inner phospholipds | scaled to 100% |
|---|---|---|---|---|
| 1 | a,e | PS 16:0-20:4 | 25.9 | 33.1 |
| 2 | f,a | PE O- 18:1-22:4 | 10.7 | 13.7 |
| 3 | a | PC 16:0-18:2 | 10.2 | 13.0 |
| 4 | a, i | PE 16:0-22:6 | 9.3 | 11.9 |
| 5 | h | PE 18:1-20:4 | 5.4 | 6.9 |
| 6 | | PC 16:0-18:1 | 5.5 | 7.0 |
| 7 | | PE 16:0-18:1 | 3.7 | 4.8 |
| 8 | g | PE O- 18:2-20:4 | 2.9 | 3.7 |
| 9 | a | PC 16:0-16:0 | 2.3 | 2.9 |
| 11 | | SM 34:1;2 | 1.5 | 2.0 |
| 10 | | PI 18:0-20:4 | 0.8 | 1.0 |

| | |
|---|---|
| a | Generally, combining lipids differing by 2 carbons and naming them by the more abundant one (eg SM 34:1 is PSM and represents this species and SM 36:1) |
| c | SMs usually have a trans double bond in the long-chain base. This is not counted as an unsaturation. This class includes SMs with an unsaturated acyl chain, including both 24:2 and shorter unsaturated spies like SM 36:2 |
| d | includes all PC species with two unsaturated acyl chains |
| e | includes lipids with the designed headgroup and one highly polyunsaturated acyl chain like 20:4 or 22:6 |
| f | For most ether lipids, there is a double bond in the vinyl ether connecting the backbone to the sn-1 hydrophobic chain. This double bond is not counted as an acyl chain unsaturation per se. Thus, this class is separate from the class which includes the more typical w-9 unsaturated acyl chains in the sn-1 (eg PE-O 18:2/20:4) |
| g | plasmalogen with C18:1 in the SN1 |

| | |
|---|---|
| h | includes all PE with both acyl chains unsaturated |
| i | includes all PE containing one saturated acyl chain and one with 2 or more unsaturations |


1 Rousselet, A. *et al.* Study of the transverse diffusion of spin labeled phospholipids in biological membranes. I. Human red bloods cells. *Biochim Biophys Acta* **426**, 357-371, (1976).

2 Verkleij, A. J. *et al.* The asymmetric distribution of phospholipids in the human red cell membrane. A combined study using phospholipases and freeze-etch electron microscopy. *Biochim Biophys Acta* **323**, 178-193, (1973).

3 Boon, J. M. & Smith, B. D. Chemical control of phospholipid distribution across bilayer membranes. *Medicinal research reviews* **22**, 251-281, (2002).

4 Bretscher, M. S. Asymmetrical lipid bilayer structure for biological membranes. *Nature: New biology* **236**, 11-12, (1972).

5 Veatch, S. L. & Keller, S. L. Separation of liquid phases in giant vesicles of ternary mixtures of phospholipids and cholesterol. *Biophys J* **85**, 3074-3083, (2003).

6 Korlach, J., Schwille, P., Webb, W. W. & Feigenson, G. W. Characterization of lipid bilayer phases by confocal microscopy and fluorescence correlation spectroscopy. *Proceedings of the National Academy of Sciences* **96**, 8461, (1999).

7 Heberle, F. A. & Feigenson, G. W. Phase separation in lipid membranes. *Cold Spring Harbor perspectives in biology* **3**, (2011).

8 Owen, D. M. *et al.* Quantitative imaging of membrane lipid order in cells and organisms. *Nat Protoc* **7**, 24-35, (2011).

9 Sezgin, E., Sadowski, T. & Simons, K. Measuring lipid packing of model and cellular membranes with environment sensitive probes. *Langmuir* **30**, 8160-8166, (2014).

10 Sezgin, E. *et al.* Elucidating membrane structure and protein behavior using giant plasma membrane vesicles. *Nat Protoc* **7**, 1042-1051, (2012).

11 Owen, D. M. *et al.* Fluorescence lifetime imaging provides enhanced contrast when imaging the phase-sensitive dye di-4-ANEPPDHQ in model membranes and live cells. *Biophys J* **90**, L80-82, (2006).

12 McIntyre, J. C. & Sleight, R. G. Fluorescence assay for phospholipid membrane asymmetry. *Biochemistry* **30**, 11819-11827, (1991).

13 Wang, Y. *et al.* Spectral characterization of the voltage-sensitive dye di-4-ANEPPDHQ applied to probing live primary and immortalized neurons. *Optics express* **17**, 984-990, (2009).

14 Levental, K. R. *et al.* omega-3 polyunsaturated fatty acids direct differentiation of the membrane phenotype in mesenchymal stem cells to potentiate osteogenesis. *Science advances* **3**, eaao1193, (2017).

15 Levental, K. R. *et al.* Polyunsaturated lipids regulate membrane domain stability by tuning membrane order. *Biophys J* **110(8)**, 1800-1810, (2016).

16 Surma, M. A. *et al.* An automated shotgun lipidomics platform for high throughput, comprehensive, and quantitative analysis of blood plasma intact lipids. *European journal of lipid science and technology : EJLST* **117**, 1540-1549, (2015).

17 Sampaio, J. L. *et al.* Membrane lipidome of an epithelial cell line. *Proc Natl Acad Sci U S A* **108**, 1903-1907, (2011).

18 Ejsing, C. S. *et al.* Global analysis of the yeast lipidome by quantitative shotgun mass spectrometry. *Proc Natl Acad Sci U S A* **106**, 2136-2141, (2009).

19 Gerl, M. J. *et al.* Quantitative analysis of the lipidomes of the influenza virus envelope and MDCK cell apical membrane. *J Cell Biol* **196**, 213-221, (2012).

20 Liebisch, G. *et al.* High throughput quantification of cholesterol and cholesteryl ester by electrospray ionization tandem mass spectrometry (ESI-MS/MS). *Biochim Biophys Acta* **1761**, 121-128, (2006).

21 Wu, E. L. *et al.* CHARMM-GUI Membrane Builder toward realistic biological membrane simulations. *Journal of Computational Chemistry* **35**, 1997-2004, (2014).

22 Klauda, J. B. *et al.* Update of the CHARMM All-Atom Additive Force Field for Lipids: Validation on Six Lipid Types. *The Journal of Physical Chemistry B* **114**, 7830-7843, (2010).

23 Venable, Richard M. *et al.* CHARMM All-Atom Additive Force Field for Sphingomyelin: Elucidation of Hydrogen Bonding and of Positive Curvature. *Biophysical Journal* **107**, 134-145, (2014).

24 Phillips, J. C. *et al.* Scalable molecular dynamics with NAMD. *Journal of Computational Chemistry* **26**, 1781-1802, (2005).

25 Feller, S. E., Zhang, Y., Pastor, R. W. & Brooks, B. R. Constant pressure molecular dynamics simulation: The Langevin piston method. *The Journal of Chemical Physics* **103**, 4613-4621, (1995).
26 Martyna, G. J., Tobias, D. J. & Klein, M. L. Constant pressure molecular dynamics algorithms. *The Journal of Chemical Physics* **101**, 4177-4189, (1994).
27 Essmann, U. *et al.* A smooth particle mesh Ewald method. *The Journal of Chemical Physics* **103**, 8577-8593, (1995).
28 Shaw, D. E. *et al.* in *SC '14: Proceedings of the International Conference for High Performance Computing, Networking, Storage and Analysis.* 41-53.
29 Hoover, W. G. Canonical dynamics: Equilibrium phase-space distributions. *Physical Review A* **31**, 1695-1697, (1985).
30 Nosé, S. A unified formulation of the constant temperature molecular dynamics methods. *The Journal of Chemical Physics* **81**, 511-519, (1984).
31 Shan, Y. *et al.* Gaussian split Ewald: A fast Ewald mesh method for molecular simulation. *The Journal of Chemical Physics* **122**, 054101, (2005).
32 Beaven, A. H. *et al.* Gramicidin A Channel Formation Induces Local Lipid Redistribution I: Experiment and Simulation. *Biophysical Journal* **112**, 1185-1197, (2017).
33 Camley, B. A., Lerner, M. G., Pastor, R. W. & Brown, F. L. H. Strong influence of periodic boundary conditions on lateral diffusion in lipid bilayer membranes. *The Journal of Chemical Physics* **143**, 243113, (2015).
34 Cui, H., Lyman, E. & Voth, G. A. Mechanism of Membrane Curvature Sensing by Amphipathic Helix Containing Proteins. *Biophysical journal* **100**, 1271-1279, (2011).
35 Diaz-Rohrer, B. B., Levental, K. R., Simons, K. & Levental, I. Membrane raft association is a determinant of plasma membrane localization. *Proc Natl Acad Sci U S A* **111**, 8500-8505, (2014).
36 Levental, I. *et al.* Palmitoylation regulates raft affinity for the majority of integral raft proteins. *Proc Natl Acad Sci U S A* **107**, 22050-22054, (2010).
37 Lorent, J. H. *et al.* Structural determinants and functional consequences of protein affinity for membrane rafts. *Nature communications* **8**, 1219, (2017).
38 Li, Q. *et al.* Electroformation of giant unilamellar vesicles in saline solution. *Colloids and surfaces. B, Biointerfaces* **147**, 368-375, (2016).
39 Steinkuhler, J. *et al.* Charged giant unilamellar vesicles prepared by electroformation exhibit nanotubes and transbilayer lipid asymmetry. *Scientific reports* **8**, 11838, (2018).
40 Horger, K. S., Estes, D. J., Capone, R. & Mayer, M. Films of agarose enable rapid formation of giant liposomes in solutions of physiologic ionic strength. *J Am Chem Soc* **131**, 1810-1819, (2009).
41 Murphy, J. & Riley, J. P. A modified single solution method for the determination of phosphate in natural waters. *Analytica Chimica Acta* **27**, 31-36, (1962).
42 Yuan, Z. *et al.* Predicting the solvent accessibility of transmembrane residues from protein sequence. *J Proteome Res* **5**, 1063-1070, (2006).
43 Dinic, J., Biverstahl, H., Maler, L. & Parmryd, I. Laurdan and di-4-ANEPPDHQ do not respond to membrane-inserted peptides and are good probes for lipid packing. *Biochim Biophys Acta* **1808**, 298-306, (2011).
44 Sharpe, H. J., Stevens, T. J. & Munro, S. A comprehensive comparison of transmembrane domains reveals organelle-specific properties. *Cell* **142**, 158-169, (2010).
45 Tanaka, K. I. & Ohnishi, S. Heterogeneity in the fluidity of intact erythrocyte membrane and its homogenization upon hemolysis. *Biochim Biophys Acta* **426**, 218-231, (1976).
46 Seigneuret, M., Zachowski, A., Hermann, A. & Devaux, P. F. Asymmetric lipid fluidity in human erythrocyte membrane: new spin-label evidence. *Biochemistry* **23**, 4271-4275, (1984).
47 Cogan, U. & Schachter, D. Asymmetry of lipid dynamics in human erythrocyte membranes studied with impermeant fluorophores. *Biochemistry* **20**, 6396-6403, (1981).
48 Schroeder, F. Differences in fluidity between bilayer halves of tumour cell plasma membranes. *Nature* **276**, 528-530, (1978).
49 Igbavboa, U., Avdulov, N. A., Schroeder, F. & Wood, W. G. Increasing age alters transbilayer fluidity and cholesterol asymmetry in synaptic plasma membranes of mice. *Journal of neurochemistry* **66**, 1717-1725, (1996).
50 Schachter, D., Abbott, R. E., Cogan, U. & Flamm, M. Lipid fluidity of the individual hemileaflets of human erythrocyte membranes. *Annals of the New York Academy of Sciences* **414**, 19-28, (1983).
51 Nikolova-Karakashian, M. N., Petkova, H. & Koumanov, K. S. Influence of cholesterol on sphingomyelin metabolism and hemileaflet fluidity of rat liver plasma membranes. *Biochimie* **74**, 153-159, (1992).
52 Morrot, G. *et al.* Asymmetric lateral mobility of phospholipids in the human erythrocyte membrane. *Proc Natl Acad Sci U S A* **83**, 6863-6867, (1986).

53 el Hage Chahine, J. M., Cribier, S. & Devaux, P. F. Phospholipid transmembrane domains and lateral diffusion in fibroblasts. *Proc Natl Acad Sci U S A* **90**, 447-451, (1993).

54 Rimon, G., Meyerstein, N. & Henis, Y. I. Lateral mobility of phospholipids in the external and internal leaflets of normal and hereditary spherocytic human erythrocytes. *Biochim Biophys Acta* **775**, 283-290, (1984).

55 Golebiewska, U. *et al.* Diffusion coefficient of fluorescent phosphatidylinositol 4,5-bisphosphate in the plasma membrane of cells. *Mol Biol Cell* **19**, 1663-1669, (2008).

56 Lipsky, N. G. & Pagano, R. E. Sphingolipid metabolism in cultured fibroblasts: microscopic and biochemical studies employing a fluorescent ceramide analogue. *Proc Natl Acad Sci U S A* **80**, 2608-2612, (1983).
